# Robust parameter estimation and identifiability analysis with Hybrid Neural Ordinary Differential Equations in Computational Biology

**DOI:** 10.1101/2024.06.04.597372

**Authors:** Stefano Giampiccolo, Federico Reali, Anna Fochesato, Giovanni Iacca, Luca Marchetti

## Abstract

Parameter estimation is one of the central problems in computational modeling of biological systems. Typically, scientists must fully specify the mathematical structure of the model, often expressed as a system of ordinary differential equations, to estimate the parameters. This process poses significant challenges due to the necessity for a detailed understanding of the underlying biological mechanisms. In this paper, we present an approach for estimating model parameters and assessing their identifiability in situations where only partial knowledge of the system structure is available. The partially known model is extended into a system of Hybrid Neural Ordinary Differential Equations, which captures the unknown portions of the system using neural networks.

Integrating neural networks into the model structure introduces two primary challenges for parameter estimation: the need to globally explore the search space while employing gradient-based optimization, and the assessment of parameter identifiability, which may be hindered by the expressive nature of neural networks. To overcome the first issue, we treat biological parameters as hyperparameters in the extended model, exploring the parameter search space during hyperparameter tuning. The second issue is then addressed by an *a posteriori* analysis of parameter identifiability, computed by introducing a variant of a well-established approach for mechanistic models. These two components are integrated into an end-to-end pipeline that is thoroughly described in the paper. We assess the effectiveness of the proposed workflow on test cases derived from three different benchmark models. These test cases have been designed to mimic real-world conditions, including the presence of noise in the training data and various levels of data availability for the system variables.

**Author summary:** Parameter estimation is a central challenge in modeling biological systems. Typically, scientists calibrate the parameters by aligning model predictions with measured data once the model structure is defined. Our paper introduces a workflow that leverages the integration between mechanistic modeling and machine learning to estimate model parameters when the model structure is not fully known. We focus mainly on analyzing the identifiability of the model parameters, which measures how confident we can be in the parameter estimates given the available experimental data and partial mechanistic understanding of the system. We assessed the effectiveness of our approach in various *in silico* scenarios. Our workflow represents a first step to adapting traditional methods used in fully mechanistic models to the scenario of hybrid modeling.

## Introduction

Mathematical and computational models are increasingly employed in the study of biological systems [1]. These models not only facilitate the creation of predictive and explanatory tools [2] but also offer a means to understand the interactions among the variables of the system [3, 4].

The well-established approach to develop a mathematical model for biological systems, *i*.*e*., the mechanistic modeling approach [5], relies on encoding known biological mechanisms into systems of partial or ordinary differential equations (PDE or ODE) using suitable kinetic laws, *e*.*g*., the law of mass action or Michaelis-Menten kinetics [6]. These equations typically incorporate several unknown model parameters, making parameter estimation a central challenge in model development [7]. The prevailing approach to parameter estimation involves optimizing the model dynamics to align with experimental data [8, 9]. Various optimization techniques are employed for this purpose, including linear and nonlinear least squares methods [10, 11], genetic and evolutionary algorithms [9], Bayesian Optimization [12, 13], control theory-derived approaches [8, 14] and, more recently, physics-informed neural networks [7]. The scarcity of experimental data and measurement noise often leads to non-identifiability issues, where the optimization problem lacks a unique solution [15]. To address this challenge, various methods exist to analyze the identifiability of model parameters. *Structural identifiability* analysis [16], performed before parameter estimation, analyzes the structure of the model to determine if parameters can be uniquely estimated. On the other hand, *practical identifiability* analysis [17], conducted after parameter estimation, evaluates how uncertainties in experimental measurements affect parameter estimation.

Developing mathematical models and estimating model parameters with the mechanistic modeling approach presents significant challenges due to the need for a detailed understanding of the interactions between (and within) biological systems. Complex biological systems often involve processes at different scales, such as genetic, molecular, tissue, organ, or whole-body, and such intricate mechanistic details are only partially known [18, 19]. To overcome the limits of mechanistic modeling, hybrid models that combine mechanistic ODE-based dynamics with neural network components have grown in popularity in various scientific domains [20, 21]. The integration between neural networks and ODE systems is known by different names, such as *Hybrid Neural Ordinary Differential Equations* (HNODEs) [22], *graybox modeling* [23, 24], or *universal differential equations* [25]. In a HNODE, neural networks are used as universal approximators to represent unknown portions of the system. Mathematically, HNODE can be formulated as the following ODE:

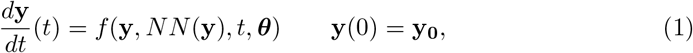

where *NN* denotes the neural network component of the model, *f* encodes the mechanistic knowledge of the system, and ***θ*** represents a vector of unknown mechanistic parameters. This approach has proven successful in several fields, and early results in isolated and relatively simple scenarios have shown promise in computational biology as well [26, 27]. However, modeling with HNODEs remains an active area of research, and best practices for estimating mechanistic parameters and assessing their identifiability within this framework remain to be outlined.

Mechanistic parameter estimation within the HNODE framework presents significant challenges. Firstly, while model calibration in mechanistic models usually relies on global optimization techniques to explore the parameter search space [11], training HNODE models necessitates the use of local and gradient-based methods [25]. Secondly, incorporating a universal approximator, such as a neural network, into a dynamical model may compromise the identifiability of the HNODE mechanistic components [28, 29]. In this sense, the existing literature has focused on trying to enforce the identifiability of the mechanistic parameters within a HNODE by integrating a regularization term into the cost function [30]. Common choices for this regularization term include minimizing the impact of the neural network on the model [30] or ensuring that the outputs of the neural network and mechanistic part are uncorrelated [28]. However, these approaches do not always guarantee the correct identification of the mechanistic part, and the outcomes depend on the specific regularization term used [28]. To the best of our knowledge, the identifiability analysis of the mechanistic parameters in a HNODE model has not been investigated in the literature so far.

In this contribution, we present an end-to-end approach for mechanistic parameter estimation and identifiability analysis in scenarios where mechanistic knowledge about the system is incomplete. Initially, we focus on tuning hyperparameters to embed our incomplete mechanistic model into a HNODE model that is capable of effectively capturing the experimental dynamics. Subsequently, we proceed to compute mechanistic parameter estimates by training the HNODE model. Following parameter estimation, we extend a well-established approach for mechanistic models to assess the parameter identifiability *a posteriori*. For identifiable parameters, we finally estimate asymptotic confidence intervals (CIs).

The proposed approach has been tested in three different *in silico* scenarios, that have been constructed by assuming a lack of information about some portions of known mechanistic models. We aimed to replicate typical conditions found in real-world scenarios, with a noisy and scattered training set describing the time evolution of a subset of the model variables. Firstly, we consider the traditional Lotka-Volterra model for predator-prey interactions. Despite being a relatively simple model, it shows the ability of the approach to identify compensations between the neural network and the mechanistic component of the HNODE model. Secondly, we evaluate the performances of our approach on a model for cell apoptosis [31], known for the non-identifiability of part of its parameters. Thirdly, we consider a model of oscillations in yeast glycolysis [32], which has been frequently employed as a benchmark for inference in computational systems biology due to its nonlinear oscillatory dynamics.

## Materials and methods

A schematic representation of the workflow is presented in Fig 1. The input of the pipeline consists of an incomplete mechanistic model containing parameters to be estimated, along with a time series dataset containing experimental observations of some of the system variables. By embedding the incomplete model into a HNODE model, our approach enables parameter estimation and identifiability analysis.

**Fig 1.**
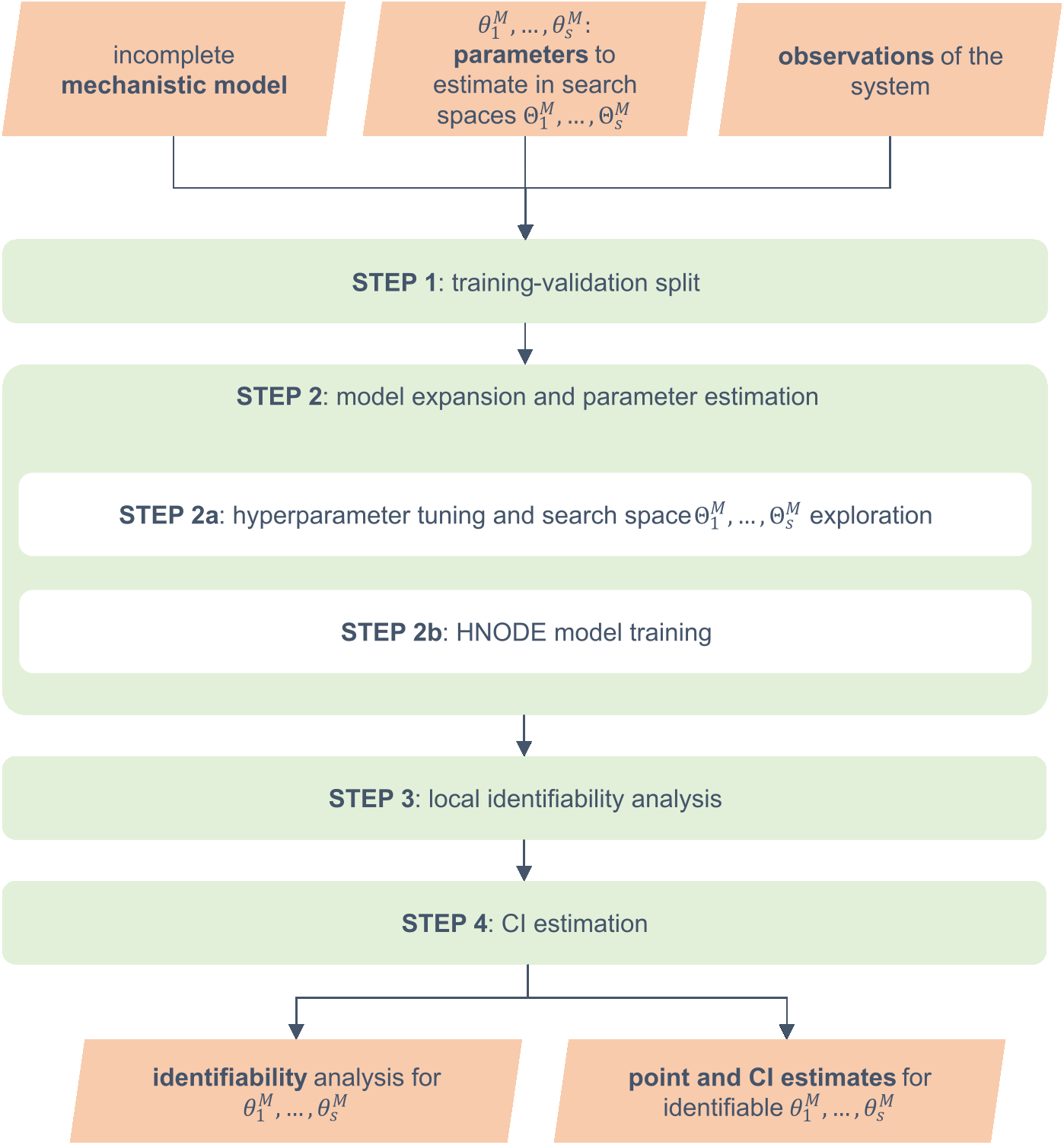
Schematic representation of the workflow. In the schema, 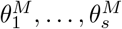 denote the mechanistic parameters to estimate in the corresponding search spaces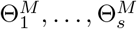 . HNODE: Hybrid Neural Ordinary Differential Equation, CI: Confidence Interval.

The workflow starts by splitting the observation time points into training and validation sets (Step 1). In the second step, using this partition, we expand the incomplete mechanistic model into a HNODE model. We employ Bayesian Optimization to simultaneously tune the model hyperparameters and explore the mechanistic parameter search space (Step 2a). The model is then fully trained (Step 2b), yielding mechanistic parameter estimates. In the next step (Step 3), we assess the local identifiability at-a-point of the parameters. For the locally identifiable ones, we proceed to estimate confidence intervals (Step 4).

In the rest of the section, we use the following notation. Let **y**(*t*, ***θ***^*M*^ , ***θ***^*NN*^ , **y**_**0**_) ∈ ℝ^*n*^, with *n* ∈ ℕ^+^, be the HNODE model defined by the following differential equation:

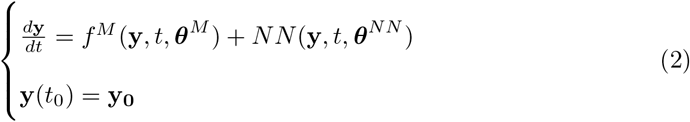

where **y**_**0**_ ∈ ℝ^*n*^ stands for the initial conditions, *f* ^*M*^ and 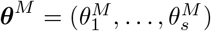 denote the mechanistic part of the model and the *s* ∈ ℕ^+^ mechanistic parameters to be estimated, respectively, while ***θ***^*NN*^ represents the neural network parameters. We denote the *i*-th component of the model as *y*^*i*^(*t*, ***θ***^*M*^ , ***θ***^*NN*^ , **y**_**0**_), with *i* = 1, … , *n*. Typically, only a subset of the system variables is observable; we indicate this subset with *O* ⊆ {1, … , *n*}. We assume to have access to the experimental measurements of the observable variables at *m* + 1 time points *t*_0_, … , *t*_*m*_; 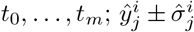 indicates the observations of the *i*-th variable at time *t*_*j*_, with the uncertainty reflecting the variability of the measurement. Additionally, we assume to have access to the initial conditions 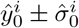 of all the variables.

In the following, we provide a comprehensive description of each step in the workflow. Given that the training of HNODE extends the training for Neural Ordinary Differential Equations (NODE) [33], the next subsection provides an overview of the methods employed for training NODE.

### Background: NODE training

In this section, to discuss the NODE case, we assume that *f*_*M*_ = **0** in Eq. (2), indicating no mechanistic knowledge about the system. Under this hypothesis, the vector field is entirely parameterized by the neural network *NN* , and the ODE:

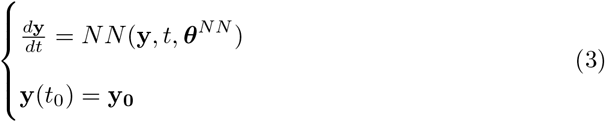

represents a NODE. Given a loss function ℒ, the training of the NODE can be achieved through the minimization of the cost function:

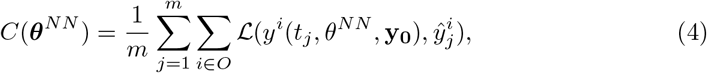

where *y*^*i*^ is computed numerically by integrating Eq. (3). Minimizing *C*(***θ***^*NN*^ ) requires to back-propagate the error through the ODE solver algorithm used for the numerical integration of *y*. By the chain rule, for a differentiable ℒ, this amounts to compute the gradients

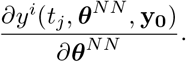

The authors in [33] demonstrated that these gradients can be efficiently computed using adjoint sensitivity [34, 35], treating the ODE solver as a black box. Adjoint sensitivity requires a backward integration of the system and different numerical methods have been proposed to efficiently calculate it [25].

As in the traditional ODE case, stiffness constitutes a significant challenge in the training of NODE [36]. However, as shown in [37], there are specific ways to overcome this issue. These include employing deep neural network architectures, *ad hoc* methods for computing adjoint sensitivity, and a normalized loss function.

### Step 1: training-validation split

In the first step of the workflow, the observation time points are divided into training and validation sets, indicated with {*t*_*j*_}_*j∈T*_ and {*t*_*j*_}_*j∈V*_ respectively. The validation time points are chosen from *t*_1_, … , *t*_*m*_ to ensure a homogeneous distribution along the observed trajectory.

### Step 2: model expansion and parameter estimation

In this step, the incompletely specified mechanistic model is expanded into a HNODE model through the use of neural networks to replace unknown portions of the system. The estimates for mechanistic parameters are derived by training the HNODE model.

The training of a HNODE model is analogous to that of NODE. Adopting a loss function ℒ, the most straightforward approach would be minimizing the cost function:

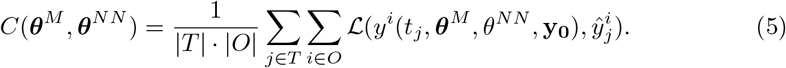

Here, |*T* | and |*O*| denote the cardinality of *T* and |*O*| respectively. Gradients of *C*(***θ***^*M*^ , ***θ***^*NN*^ ) with respect to mechanistic and neural network parameters are calculated via adjoint sensitivity methods. If the cost function defined by Eq. (5) is employed, a single trajectory of **y** spanning from *t*_0_ to *t*_*m*_ is computed at each epoch of the training. This approach, known as single shooting, is often suboptimal due to the potential risks of the training getting stuck in local minima [29].

Therefore, we adopt the multiple shooting technique (MS) [38]. In MS, the time interval (*t*_0_, *t*_*m*_) is partitioned into different segments. Rather than computing a single trajectory of **y** on the entire training interval, a potentially discontinuous trajectory **y**_*P* *W*_ (*t*, ***θ***^*M*^ , ***θ***^*NN*^ ) is reconstructed piece-wise by solving an initial value problem on each segment, adding the states of the system at the initial points of each segment as additional parameters to be optimized. The cost function is then composed of two terms:

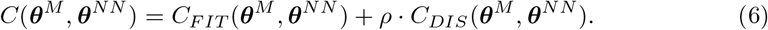

In Eq. (6), *C*_*FIT*_ measures the difference between the piecewise-defined trajectory and the training data, *C*_*DIS*_ measures the discontinuity of the trajectory, and *ρ ∈* ℝ is a hyperparameter. For *C*_*FIT*_ , we use the cost function defined in Eq. (5), adopting the loss function

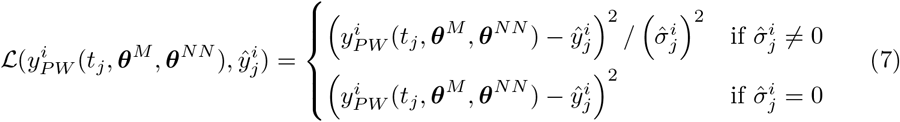

in which the quadratic loss is weighted for the uncertainty measure if it is not zero. *C*_*DIS*_ is computed by summing the squared values of the discontinuities at the extremes of the training interval partition. To prevent overfitting, in the case of noisy training data, we add an L2 regularization term. Given this cost function, the training is performed with a gradient-based optimizer, and gradients are computed using adjoint sensitivity methods. In our test cases, we use the Adam optimizer [39], and we refine the results with L-BFGS [40] on training datasets without noise.

To define the segmentation of the time interval, we partition the training set of time points into subsets containing the same number *k* of items (excluding the last interval), where *k* is a hyperparameter. When using MS to train the model, we introduce as additional parameters to optimize the state of the system at the initial points of each segment. These parameters are initialized with the measured states of the system at the time point when the corresponding system variable is observable; otherwise, they are initialized with the initial state of the variable.

### How to tune the hyperparameters and explore the parameter search space

The hyperparameters requiring tuning can be grouped into three different macro-categories: the hyperparameters of the architecture of the neural network, the starting values of the mechanistic parameters to initiate the gradient-based optimization, and the hyperparameters related to the MS training. This latter category includes the segmentation of the time interval (*t*_0_, *t*_*m*_) used to define the piecewise trajectory, the weight factor *ρ* used in Eq. (6), the weight *λ* of the *L*2 regularization term, and the learning rate of the gradient-based method. Incorporating the initial values of 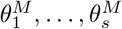 into hyperparameter tuning enables the integration of local gradient-based search with Bayesian Optimization techniques, as in [41]. The objective is to globally explore the mechanistic parameter search space during this step.

The hyperparameters are tuned in two stages, described below.

### Hyperparameter tuning - Stage 1

We simultaneously tune all hyperparameters, except for the *L*2 regularization factor *λ*, using the multivariate Tree-Structured Parzen Estimator (TPE) [42, 43]. In each TPE trial, the HNODE model is trained for a limited number of MS iterations. The optimization metric during the TPE algorithm is the mean squared error on the validation time points of the trajectory predicted by the trained model:

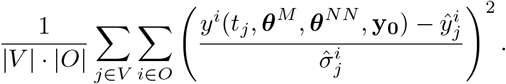

As before, we employ a pure quadratic error when uncertainty indications on measurements are absent.

### Hyperparameter tuning - Stage 2

In the second stage, we fix the hyperparameters tuned in the first step and focus on tuning the *L*2 regularization factor, *λ*, using a grid-search approach. In each trial during this step, the HNODE model is trained for an extended number of MS iterations, and we continue to employ the same metric as in the previous step to evaluate each trial.

The decision to split the tuning process into two steps is motivated by the observation that the effects of overfitting may not become apparent with a low number of iterations. Therefore, tuning *λ* in the initial stage might not be as effective. Conversely, conducting a large number of MS iterations in the first stage would significantly increase computational costs.

### How to deal with stiffness

Before starting the hyperparameter tuning phase, the behavior of the HNODE model, whether stiff or non-stiff, is unknown. To train a stiff HNODE model, we follow the guidelines outlined for stiff NODE. This involves employing specialized techniques such as *ad hoc* adjoint sensitivity methods, deeper neural network architectures, stiff ODE solvers, and normalized loss functions. We do not include all configurations used for stiff systems as hyperparameters to tune, to prevent an excessive increase in the search space dimension. Instead, we propose initiating the hyperparameter tuning with non-stiff configurations and monitoring the integration of the HNODE model during the first trials of the process, such as observing the number of steps taken by the ODE solver to complete system integration. If stiffness is detected, we finalize the tuning process and we train the HNODE model using stiff configurations.

### Step 3: local identifiability analysis

In this step, we investigate *a posteriori* the local identifiability of the mechanistic parameters 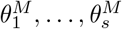 estimated with the model training. This is particularly important in HNODE models, as incorporating a universal approximator, such as a neural network, into a dynamical model may compromise the identifiability of mechanistic parameters. The following subsections introduce the foundational concepts for our approach and the description of the workflow.

### Local identifiability at-a-point and sloppy directions

We refer to the combined neural network and mechanistic parameter estimates, obtained through the training in Step 2, as the single vector:

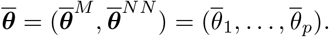

We recall that the *k*-th parameter of the model is *locally identifiable at-a-point* in 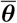[44] if

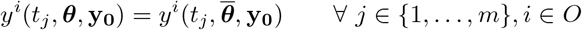

implies 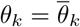 for all ***θ*** in a suitable neighborhood of 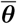.

To study the local identifiability, we quantify the change in the HNODE model behavior, as parameters ***θ*** vary from 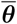, with the function:

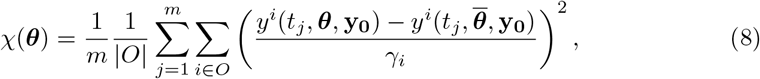

where 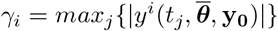 is introduced to normalize the contribution to *χ* of each variable. In a neighborhood of 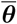, the function *χ* can be approximated as:

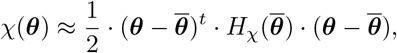

where 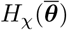 denotes the Hessian matrix of *χ* in 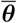. If **v** is an eigenvector of 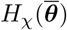 and *μ* is the corresponding eigenvalue, for small values of *α* it holds:

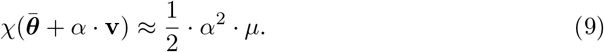

Eq. (9) implies that the HNODE model behavior does not change significantly when the parameters move along the direction identified by **v** if *μ* is zero. In mechanistic models, the eigenvectors related to zero or almost-zero eigenvalues are commonly called sloppy directions [45–47]. Denoting with *V*_0_ the null space of *H*_*χ*_, spanned by the eigenvectors related to zero eigenvalues, and with *V*_S_ the subspace spanned by the remaining eigenvectors, each parameter *θ*_*k*_ of the model can be decomposed as:

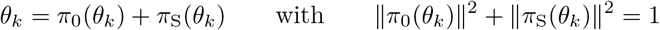

where *π*_0_(*θ*_*k*_) and *π*_S_(*θ*_*k*_) denote the projection of *θ*_*k*_ onto *V*_0_ and *V*_S_ respectively. *π*_0_(*θ*_*k*_) represents the local direction that maximizes the parameter perturbation without affecting the model behavior. Although one might expect identifiability of *θ*_*k*_ to yield ∥*π*_0_(*θ*_*k*_)∥^2^ = 0 and ∥*π*_S_(*θ*_*k*_)∥^2^ = 1, strict equalities generally do not hold. This is because, as observed in [45], each sloppy eigenvector typically encompasses mixed components of nearly all model parameters. Hence, 0 *<* ∥*π*_0_(*θ*_*k*_)∥^2^ *<* 1 applies to all model parameters.

### How to assess the local identifiability at-a-point

Our approach involves identifying mechanistic parameters 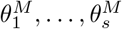 having a significant projection onto the null subspace of the HNODE model. We proceed as follows. First, we compute 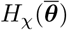 with the Gauss-Newton method:

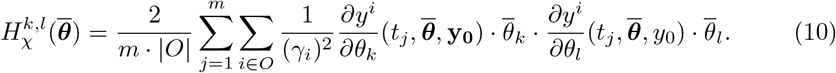

In Eq. (10), we rescale the sensitivity coefficients to make them independent of the absolute values of the parameters [48]; this consideration is important when simultaneously considering both neural network and mechanistic parameters. 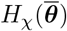 is a symmetric and positive semi-definite matrix. We proceed to determine the null subspace of 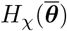 spanned by all eigenvectors corresponding to eigenvalues *μ ≤ ϵ*. Here, *ϵ* is a hyperparameter embodying the threshold used to distinguish zero eigenvalues in a numerical environment. To discriminate whether there exists a direction in the parameter search space along which we can significantly perturb the parameter 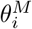 without affecting the model simulation, we analyze the norm of the projection of 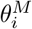 onto *V*_0_. If it exceeds a predefined threshold *δ*, which is a second hyperparameter of our approach, we classify the parameter as non-identifiable.

### Step 4: CI estimation

In the final step of our pipeline, we estimate CIs for the locally identifiable parameters. Several methods for estimating parameter CIs in mechanistic models have been proposed. We employ the Fisher Information Matrix (FIM)-based approach [49]. We assume that our observed data follow independent Gaussian distributions 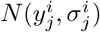, and that our parameter estimator is an approximation of the maximum likelihood estimator, as done in [7]. Therefore, we can approximate the observed FIM as:

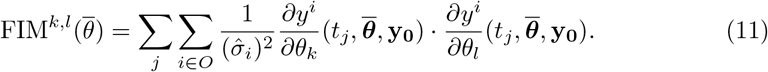

The pseudo-inverse of the FIM provides a lower bound for the covariance matrix of our estimators [50]. Consequently, we can define a lower bound 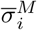 for the standard deviation of the estimator of the parameter 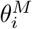 by taking the square roots of the elements on the diagonal of FIM^−1^. We can then compute the lower bound for the *α*-level confidence intervals of 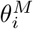 as:

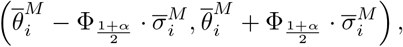

where Φ_*x*_ represents the *x* quantile of a standard Gaussian distribution.

### Algorithmic setup

All the results presented in the Results Section have been obtained employing the following hyperparameters. The observation data point datasets are partitioned into training and validation sets with an 8:2 ratio. The first stage of the hyperparameter tuning consists in 500 iterations of TPE: in each trial, the HNODE model is trained for 500 epochs using Adam optimizer. In the second stage of the hyperparameter tuning (performed only on datasets with noise) the model is trained for 2000 epochs in each trial. The comprehensive training of the HNODE model encompasses 10000 epochs utilizing the Adam optimizer. Subsequent refinement involves the application of the L-BFGS algorithm until convergence, with a maximum of 5000 epochs (performed only on noiseless datasets). The optimized mechanistic parameters are clipped to the lower or upper bounds of the parameter search spaces when they exceed those limits. Each model is trained with 10 initializations of the neural network, with weights initialized using Glorot Uniform [51]. The training resulting in the lowest validation cost is then selected. The identifiability analysis is performed using the thresholds *ϵ* = 10^−5^ and *δ* = 0.05; a discussion regarding these thresholds is provided in the Discussion Section.

### Implementation

All the computations have been run on a Debian-based Linux cluster node, with two 24-core Xeon(R) CPU X5650 CPUs and 250 GB of RAM. The code is implemented in Julia v1.9.1 [52], using the SCiML environment [25], a software suite for modeling and simulation that also incorporates machine learning algorithms. The hyperparameter tuning has been performed with Optuna [53]. The scripts to reproduce the results are available at https://github.com/cosbi-research/HNODECB.

## Results

The computational pipeline introduced herein has been tested in three different *in silico* test cases, further detailed in the subsequent sections.

### Lotka-Volterra test case

The first test case is based on the Lotka-Volterra predator-prey model, which describes the temporal dynamics of two species, one preying upon the other [54]. Although it is based on a relatively simple system, this test case offers the opportunity to evaluate the effectiveness of our approach in detecting compensations between the neural network and the mechanistic parameters of the model. Additionally, it allows for a discussion on why we opt not to enforce the identifiability of the mechanistic parameters through the use of a regularizer in the cost function, as proposed in [30].

The full mechanistic model is defined by the following system of ODE:

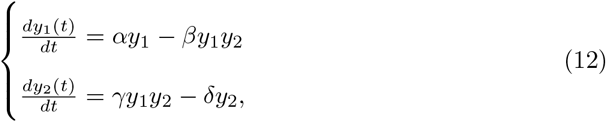

where *y*_1_ represents the prey population, and *y*_2_ represents the predator population. In this context, we assume a lack of information regarding the interaction terms between the two species (*βy*_1_*y*_2_ and *γy*_1_*y*_2_). By replacing them with a neural network *NN* : ℝ^2^ → ℝ^2^, we have the following HNODE model:

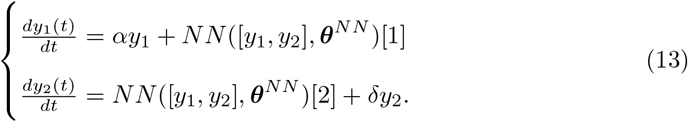

Our objective is to estimate and assess the identifiability of the prey birth rate *α*, assuming we know the predator decay rate *δ*. To generate *in silico* the observation datasets, we numerically integrate Eq. (12) over the interval from *t* = 0 to *t* = 5 years (the initial conditions and parameters are provided in Section S1 in S1 File) and sample the states every 0.25 years (resulting in 21 observed time points). We test our pipeline on a noiseless dataset (denoted with *DS*_0.00_), and on a dataset in which each time series is perturbed with a zero-mean Gaussian noise with a standard deviation equal to 5% of its min-max variation (denoted with *DS*_0.05_). The pipeline is run independently on the two datasets. Our search space for the mechanistic parameter is 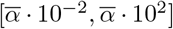, where 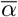 is the ground truth value used for generating the in silico observations.

We run the pipeline assuming non-stiff configurations, employing an explicit Runge-Kutta ODE solver [55] and the *Interpolating Adjoint* method [25] to compute adjoint sensitivity. Additionally, the search space for the neural network architecture is restricted to shallow networks (at most 3 hidden layers) with *tanh* as the activation function. The tuned hyperparameters, together with the search spaces, are reported in Table S3 in S1 File. The dynamics predicted by the fully trained HNODE models are illustrated in Fig 2, demonstrating the effective fitting to the observation data points. The resulting *α* estimates are reported in Table 1.

**Table 1.** Lotka-Volterra, estimated values of the prey birth rate *α*.

| Target | Dataset | Estimate |
| --- | --- | --- |
| 1.3 | $DS_{0.00}$ | 0.804 |
| | $DS_{0.05}$ | 0.744 |

**Fig 2.**
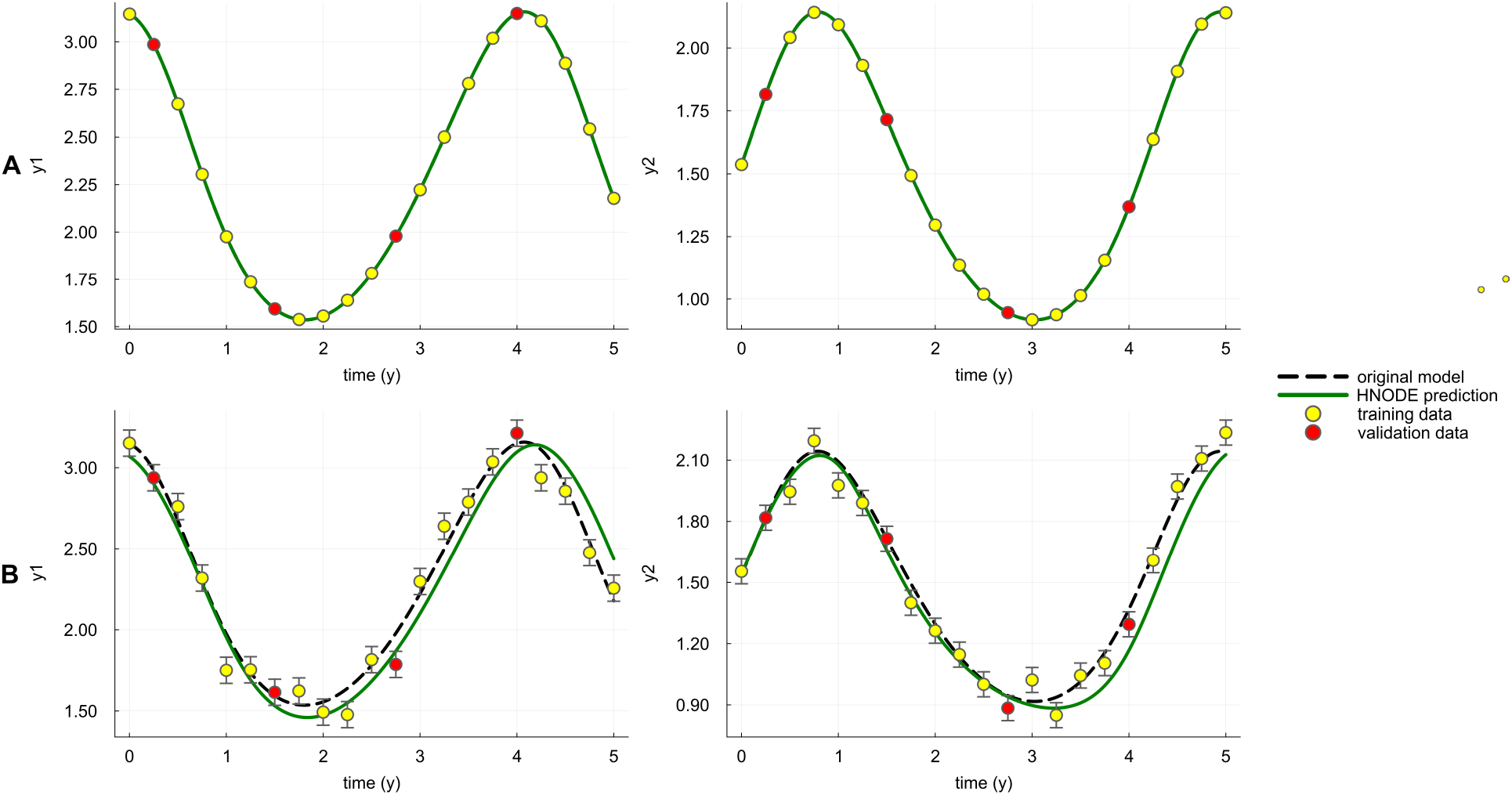
Dynamics predicted by the HNODE model in the Lotka-Volterra test case. The dynamics predicted by the HNODE model trained on *DS*_0.00_ and *DS*_0.05_ (shown in panels A and B respectively) are compared with the original model. The points represent the observations of the system, divided into training and validation sets.

To assess the identifiability of *α*, we analyze its projection onto the null subspace of *H*_*χ*_ (Fig 3). In both HNODE models, the norm of the projection exceeds *δ*, indicating the non-identifiability of the mechanistic parameter. These projections consist of both *α* and neural network parameters, suggesting a compensation between the neural network and the mechanistic parameter. This aligns with our expectations, as the first equation of Eq. (13) consists of a sum between a neural network and a mechanistic term involving *α*. It is thus plausible that the neural network, with its approximation properties, could adapt to changes in the mechanistic parameter *α*.

**Fig 3.**
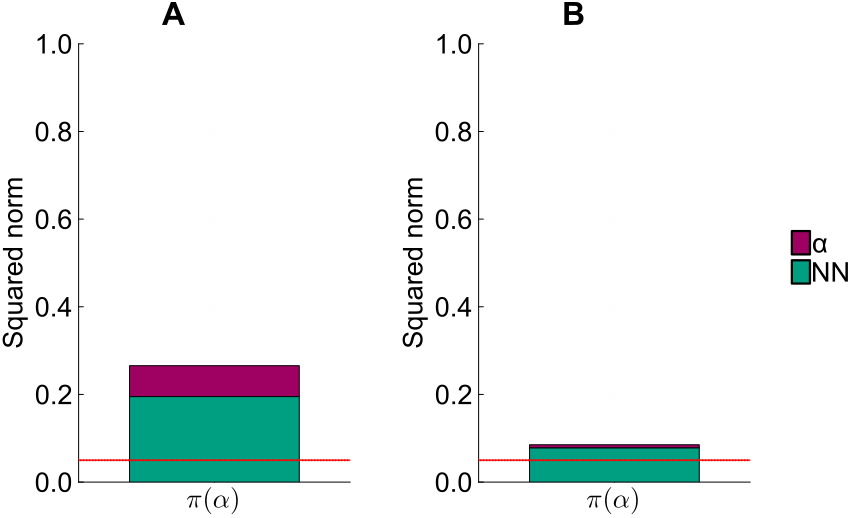
Identifiability analysis of *α* in the Lotka-Volterra test case. Squared norm of the projections of the parameter *α* onto the null subspace of *H*_*χ*_ for the models trained on *DS*_0.00_ and *DS*_0.05_ (panel A and B respectively). The total height of the bar corresponds to the squared norm of the projection, while the different components of the projection are depicted in different colors. The red line indicates the threshold to determine the identifiability of the parameter (0.05).

In this simple scenario, we aim to demonstrate qualitatively that the projection of the mechanistic parameter onto the null subspace of *H*_*χ*_ effectively embodies a compensation between the mechanistic parameter and the neural network. To achieve this, we analyze the model behavior when the parameters of the trained HNODE model are perturbed along the direction identified by the projection (Fig 4). The analysis reveals that, despite the sensitivity of the model dynamics to changes solely in the parameter *α*, moving along the projection allows a significant alteration of the value of *α* without impacting the overall model dynamics.

**Fig 4.**
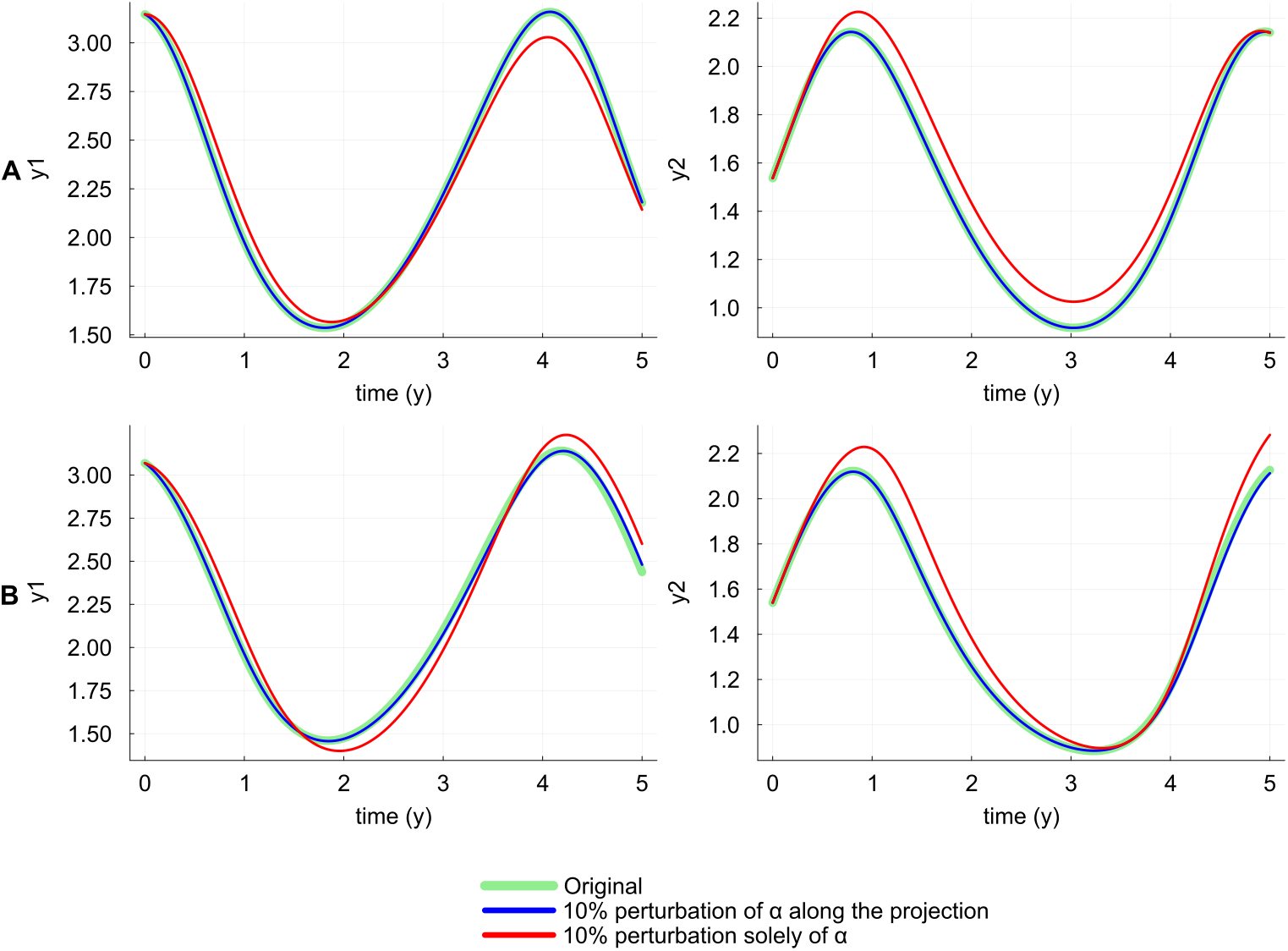
Qualitative inspection of the compensation in the Lotka-Volterra test case. The dynamics predicted by the HNODE models trained on *DS*_0.00_ and *DS*_0.05_ (panel A and B respectively) are compared with the dynamics obtained perturbing the *α* value by 10% along the projection onto the null subspace of *H*_*χ*_, and with the dynamics obtained perturbing solely *α* by 10%.

Such compensation hinders the identifiability of *α*, and our workflow terminates at this point. We conclude by showing that, in this scenario, the regularization of the cost function proposed in [30] does not ensure an accurate estimation of *α*. Their proposed regularization involves minimizing the contribution of the neural network in the HNODE model. Mathematically, this approach consists of adding the following regularizer to the cost function:

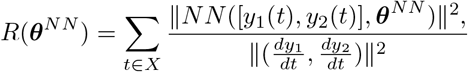

where *X* is a grid of time points on the integration interval. To demonstrate that in this case this approach would not be effective, we sampled 50 values for *α* within the interval [0.013, 3] and trained the HNODE model keeping *α* fixed. This resulted in obtaining a neural network parameterization ***θ***^*NN*^ for each *α*, yielding a profile for the regularizer:

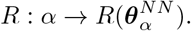

For *α ≤* 2, the value of *α* is not correlated with the ability of the HNODE model to fit the data (Fig S1 in S1 File), suggesting that the compensation holds in this region. If the inclusion of the regularizer enables the accurate estimation of *α*, the ground truth value *α* should correspond to the minimum of the regularizer profile *R*. However, our analysis indicates that this is not the case (Fig 5). Therefore, in this scenario, using the regularizer would result in a biased estimation of the parameter.

**Fig 5.**
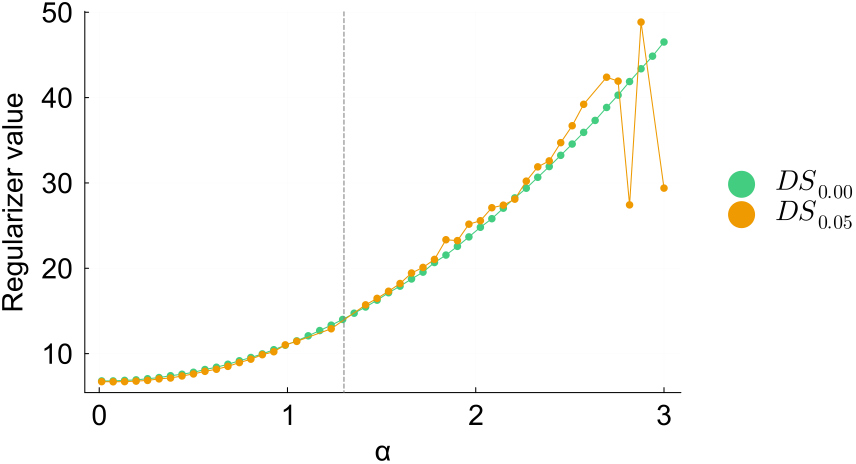
Profile of the regularizer in the Lotka-Volterra test case. The profiles of *R*(*α*) for the models trained on *DS*_0.00_ and *DS*_0.05_ are plotted in the interval [0, 3]. The vertical dashed line denotes the ground truth value of *α*.

### Cell apoptosis test case

The second test case is based on a model for cell death in apoptosis [31], which constitutes a core sub-network within the signal transduction cascade that regulates programmed cell death. This model is known for the structural and practical non-identifiability of part of its parameters [7]. The equations defining the model are as follows:

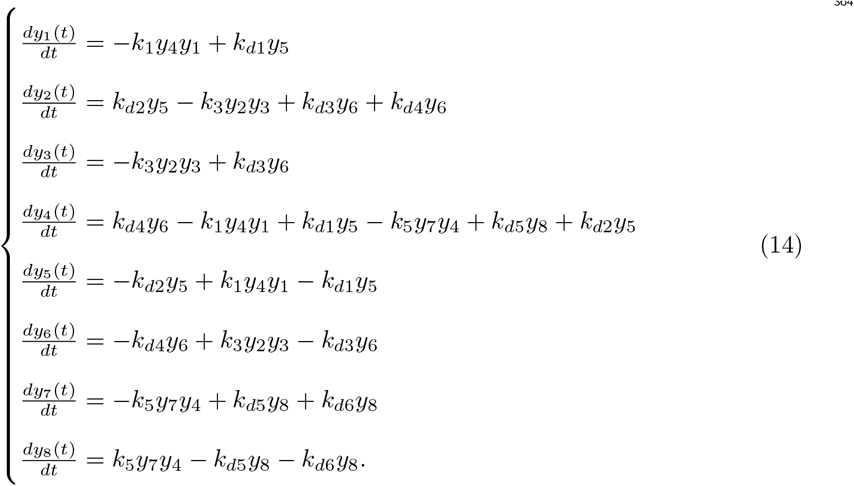

In this context, we assume a lack of mechanistic knowledge about the entire equation related to *y*_4_. We selected this variable to maximize the challenge, as the corresponding equation contains the highest number of non-linear terms in the system. Our objective is to estimate all the parameters of the model. We assume knowledge of the variables directly influencing the dynamics of *y*_4_; thus, we model the unknown derivative of *y*_4_ with the neural network:

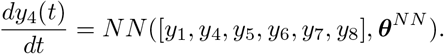

As in Lotka-Volterra, we assume a search space for the mechanistic parameters ranging from 10^−2^ to 10^2^ times the ground truth value for each parameter. These ground truth values, derived from [31], are specified in Section S5 in S1 File.

In the first scenario, we assume that all the system variables are observable. The observations are generated by numerically integrating Eq. (14) from 0 to 16 hours, with the parameters and initial conditions outlined in Section S5 in S1 File. The trajectory is sampled every 0.8 hours, resulting in 21 observed time points. Similar to the previous test case, we consider a dataset *DS*_0.00_ without noise, and a dataset *DS*_0.05_ in which each time series is perturbed with zero-mean Gaussian noise with a standard deviation equal to 5% of its min-max variation. In this test case, after observing the integration in the first trials of the hyperparameter tuning, we manually switched to stiff configurations. These configurations include utilizing the *TRBDF2* [56] implicit Runge-Kutta ODE solver and the *QuadratureAdjoint* [37] method for adjoint sensitivity computation. Additionally, the neural network architecture search spaces are expanded to include a deeper neural network (up to 6 hidden layers), with *gelu* as the activation function. The hyperparameters selected by the pipeline are reported in Table S7 in S1 File, and the dynamics for *y*_1_ and *y*_2_ predicted by the trained HNODE models are presented in Fig 6 (Figs S3 and S4 in S1 File for the dynamics of all the variables). The trained HNODE model effectively fits the data.

**Fig 6.**
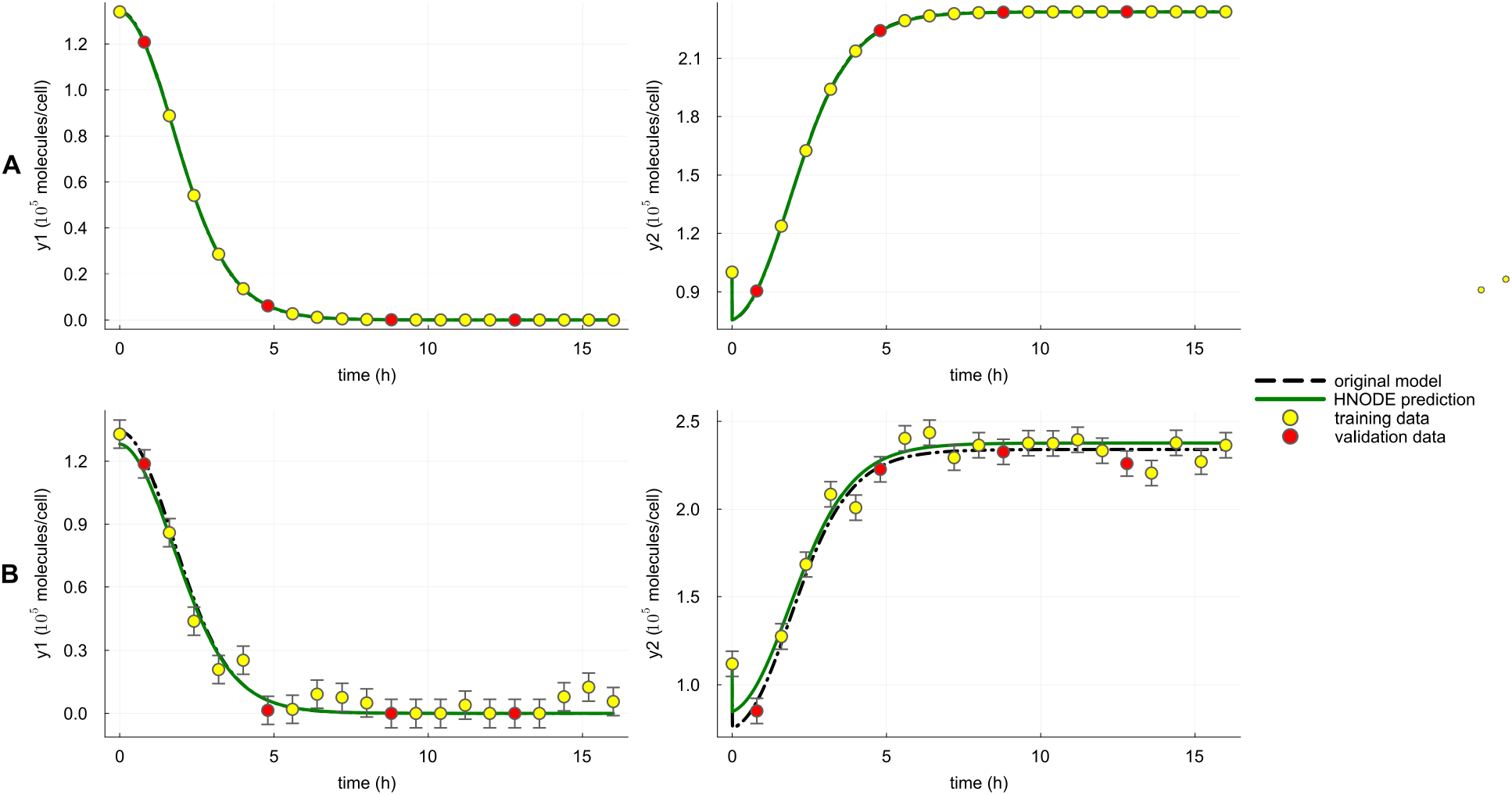
Dynamics predicted by the HNODE model in the cell apoptosis test case, first scenario. The dynamics of *y*_1_ and *y*_2_ predicted by the HNODE model trained on *DS*_0.00_ and *DS*_0.05_ (shown in panels A and B respectively) are compared with the original model. The points represent the observations of the system, divided into training and validation sets.

With the trained HNODE models, we proceed to the identifiability analysis and the CI estimations of the mechanistic parameters of the model (Table 2). Two of the model parameters, *k*_*d*2_ and *k*_*d*4_, are classified as identifiable, and this is empirically confirmed by the accuracy of their estimates (maximum relative error of 0.38% when estimated on the dataset without noise and 5.08% on the dataset with noise).

**Table 2.**
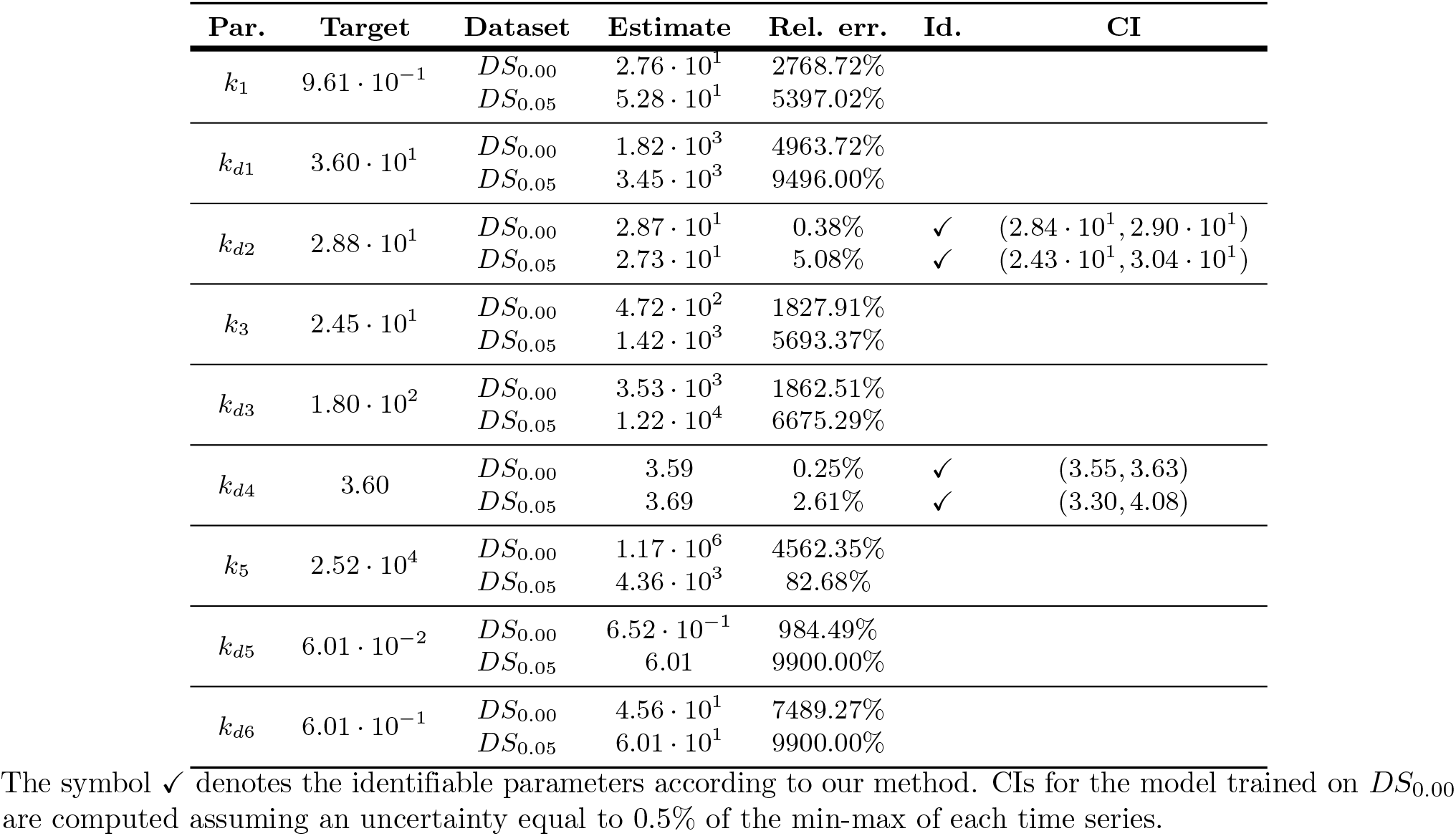
Cell apoptosis, first scenario - estimated values of mechanistic parameters.

To understand why the other mechanistic parameters are classified as not identifiable, we analyze the components of the projections on the null subspace of *H*_*χ*_. The analysis reveals the absence of neural network parameter components in both the HNODE models (Fig 7 for the model trained on *DS*_0.00_, Fig S2 in S1 File for analogous results obtained with the model trained on *DS*_0.05_). This implies that the parameter non-identifiability is not caused by compensations due to the neural network. Notably, the projections of parameters *k*_1_ and *k*_*d*1_ show almost equal contributions from both the parameters, and the same holds for the parameters *k*_3_ and *k*_*d*3_. This suggests the existence of local compensations between *k*_1_ and *k*_*d*1_, as well as between *k*_3_ and *k*_*d*3_.

**Fig 7.**
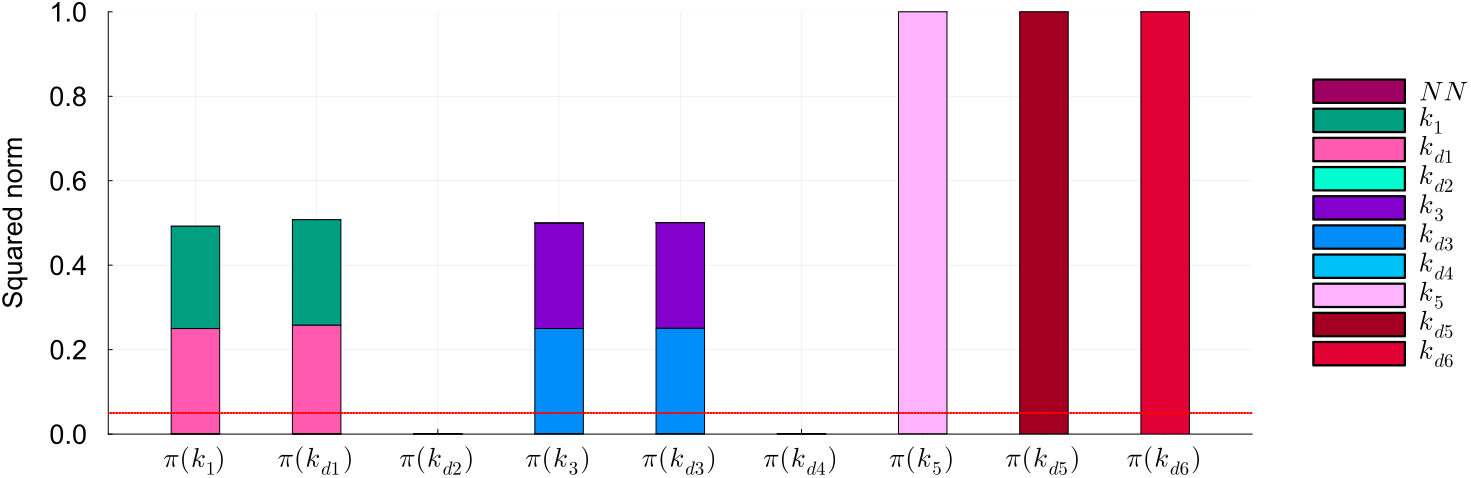
Identifiability analysis of mechanistic parameters in the cell apoptosis test case, first scenario (*DS*_0.00_). Squared norm of the projections of the mechanistic parameters onto the null subspace of *H*_*χ*_ for the model trained on *DS*_0.00_. The total height of the bar corresponds to the squared norm of the projection, while the different components of the projection are depicted in different colors. The red line indicates the threshold to determine the identifiability of the parameter (0.05).

To assess the existence of such compensations and to evaluate the performance of our approach with a larger number of identifiable parameters, we fix *k*_*d*1_ and *k*_3_ to their ground truth values (36.0 h^−1^ and 24.48cell · h^−1^ · 10^−5^ · molecules^−1^ respectively [31]) and execute the pipeline. The selected hyperparameters are listed in Table S8 in S1 File; the dynamics predicted by the HNODE models effectively fit the experimental data (Figs S6 and S7 in S1 File). The results show that in this condition the identifiability of *k*_*d*1_ and *k*_*d*3_ is restored, allowing for the estimation of four model parameters with low relative error (Table 3).

**Table 3.**
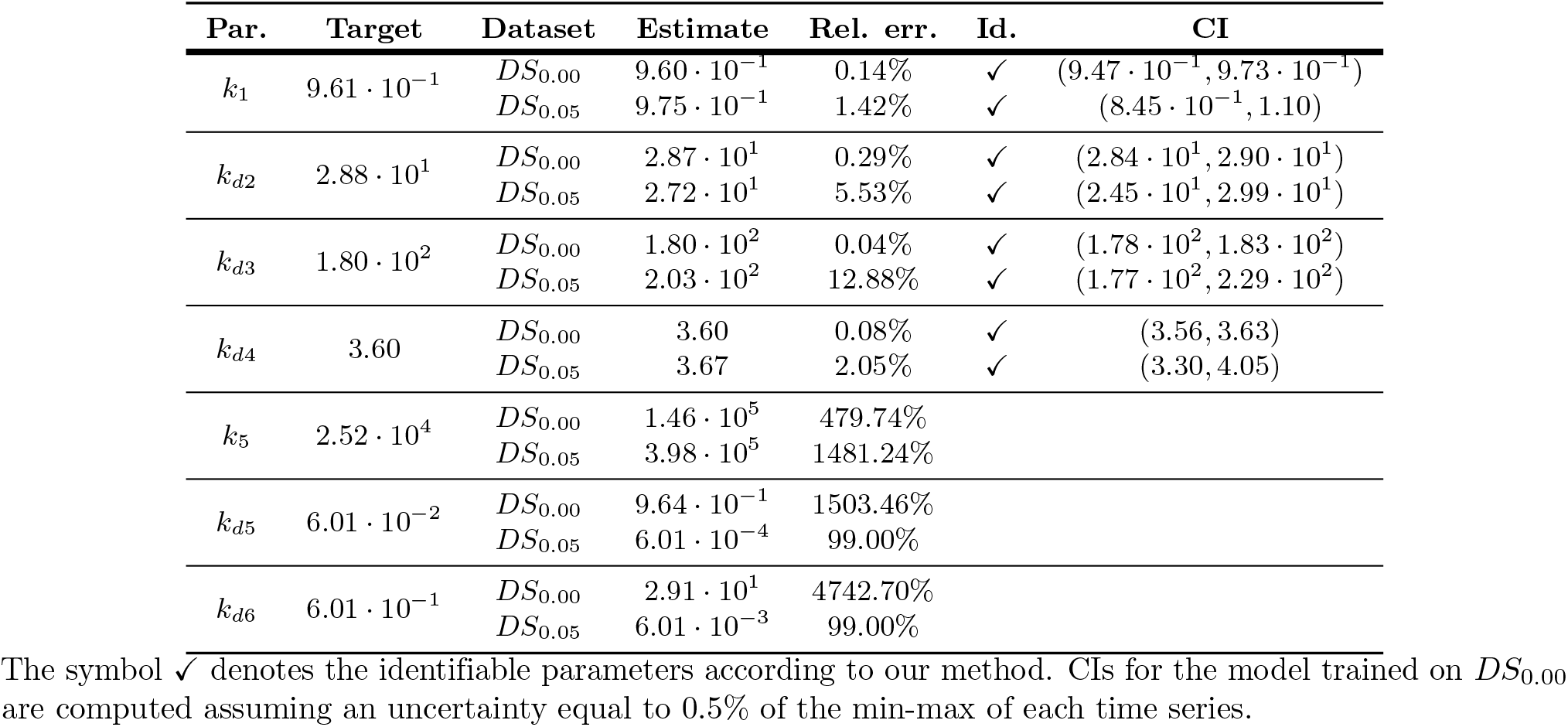
Cell apoptosis, first scenario, assuming *k*_*d*1_ and *k*_3_ fixed to their ground truth values - estimated values of mechanistic parameters.

To mimic more realistic conditions in the second scenario, keeping *k*_*d*1_ and *k*_3_ fixed to their ground truth values, we suppose that only the variables *y*_5_ and *y*_6_ are observable. We selected these variables because, within the mechanistic model, they represent the minimal subset that maximizes the number of identifiable parameters (Section S15 in S1 File). The *in silico* observations are generated as in the first scenario. However, since only 2 out of 8 model variables are observable, trajectories are sampled every 0.4 hour to maintain a comparable amount of information in the training set, resulting in 42 time points. The hyperparameters tuned in the pipeline are listed in Table S9 in S1 File, and the dynamics of the observable variables predicted by the HNODE models are shown in Figs S9 and S10 in S1 File.

In this scenario *k*_*d*2_, *k*_*d*3_, and *k*_*d*4_ are still identifiable (Table 4), and can be accurately estimated (maximum relative error of 0.17% when estimated on the dataset without noise, 11.64% on the dataset with noise), whereas *k*_1_ is not. The analysis of the components of the projections of the mechanistic parameters onto the null space of *H*_*χ*_ suggests that the non-identifiability of *k*_1_ in this scenario is caused by a compensation between *k*_1_ and the neural network (Fig 8 and Fig S8 in S1 File).

**Table 4.** Cell apoptosis, second scenario - estimated values of mechanistic parameters.

| Par. | Target | Dataset | Estimate | Rel. err. | Id. | CI |
| --- | --- | --- | --- | --- | --- | --- |
| $k_1$ | $9.61 \cdot 10^{-1}$ | $DS_{0.00}$ | 1.32 | 37.32% | | |
| | | $DS_{0.05}$ | 1.18 | 23.06% | | |
| $k_{d2}$ | $2.88 \cdot 10^1$ | $DS_{0.00}$ | $2.88 \cdot 10^1$ | 0.16% | ✓ | $(2.45 \cdot 10^1, 3.32 \cdot 10^1)$ |
| | | $DS_{0.05}$ | $2.95 \cdot 10^1$ | 2.60% | ✓ | $(-6.86 \cdot 10^1, 1.28 \cdot 10^2)$ |
| $k_{d3}$ | $1.80 \cdot 10^2$ | $DS_{0.00}$ | $1.80 \cdot 10^2$ | 0.17% | ✓ | $(1.76 \cdot 10^2, 1.85 \cdot 10^2)$ |
| | | $DS_{0.05}$ | $2.01 \cdot 10^2$ | 11.64% | ✓ | $(1.60 \cdot 10^2, 2.42 \cdot 10^2)$ |
| $k_{d4}$ | 3.60 | $DS_{0.00}$ | 3.60 | 0.03% | ✓ | (3.56, 3.64) |
| | | $DS_{0.05}$ | 3.90 | 8.22% | ✓ | (3.42, 4.37) |
| $k_5$ | $2.52 \cdot 10^4$ | $DS_{0.00}$ | $1.79 \cdot 10^6$ | 7012.35% | | |
| | | $DS_{0.05}$ | $2.62 \cdot 10^3$ | 89.59% | | |
| $k_{d5}$ | $6.01 \cdot 10^{-2}$ | $DS_{0.00}$ | $6.01 \cdot 10^{-4}$ | 99.00% | | |
| | | $DS_{0.05}$ | 4.93 | 8095.41% | | |
| $k_{d6}$ | $6.01 \cdot 10^{-1}$ | $DS_{0.00}$ | $6.01 \cdot 10^{-3}$ | 99.00% | | |
| | | $DS_{0.05}$ | $6.01 \cdot 10^1$ | 9900.00% | | |

The symbol ✓ denotes the identifiable parameters according to our method. CIs for the model trained on $DS_{0.00}$ are computed assuming an uncertainty equal to 0.5% of the min-max of each time series.

**Fig 8.**
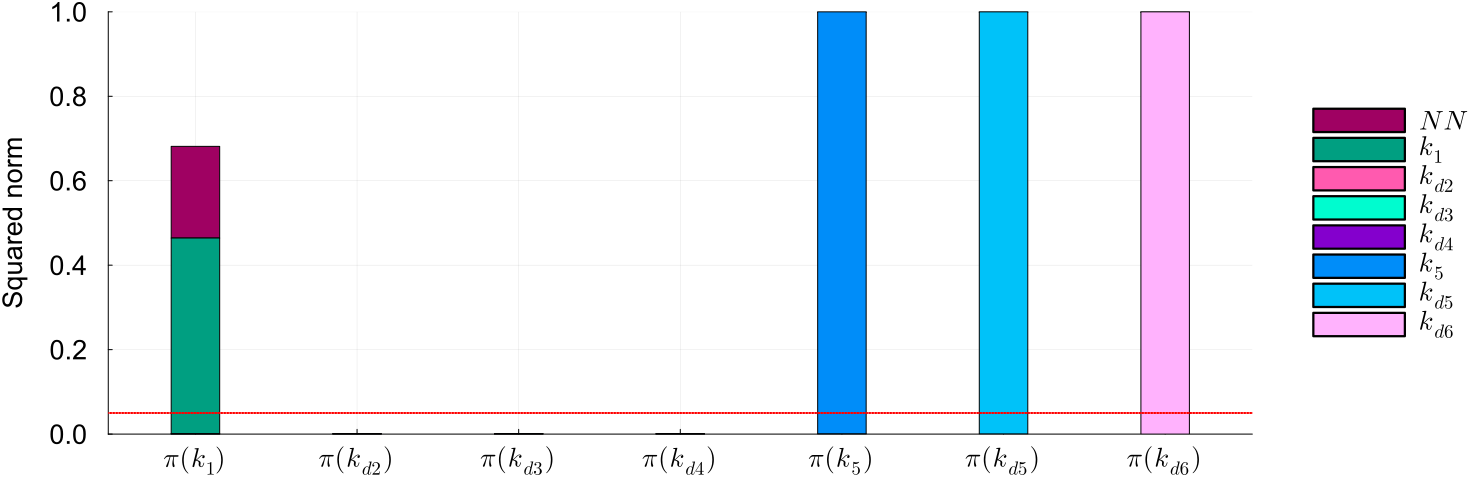
Identifiability analysis of mechanistic parameters in the cell apoptosis test case, second scenario (*DS*_0.00_). Squared norm of the projections of the mechanistic parameters onto the null subspace of *H*_*χ*_ for the model trained on *DS*_0.00_. The total height of the bar corresponds to the squared norm of the projection, while the different components of the projection are depicted in different colors. The red line indicates the threshold to determine the identifiability of the parameter (0.05).

### Yeast glycolysis model

The last test case is based on a model of oscillations in yeast glycolysis [32], which has been frequently employed as a benchmark for inference in computational systems biology [7, 57]. The model is described by the following system of ODE:

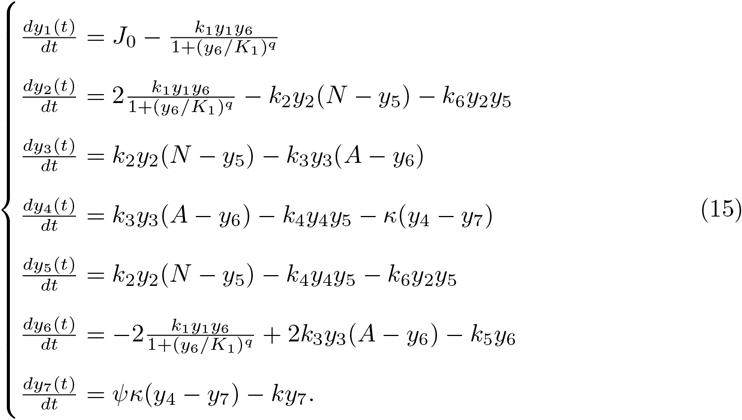

In this test case, we assume a complete absence of mechanistic knowledge concerning *y*_1_. This variable has been chosen to maximize the challenge, as its nonlinear oscillatory dynamics have proven to be particularly difficult for automatic identification of dynamical systems [58]. Our objective is to estimate all the parameters of the systems (except *J*_0_, which appears only in the derivative of *y*_1_, and therefore is not present in the HNODE model). As in the cell apoptosis case, we assume a knowledge of the variables directly influencing the dynamics of *y*_1_, thus the equation of *y*_1_ will be approximated by a neural network depending solely on *y*_1_ and *y*_6_:

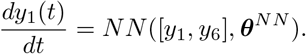

Given the greater number of system parameters compared to the first two test cases, we assume a search space ranging from 10^−1^ to 10^1^ times the ground truth value for each mechanistic parameter. These ground truth values, derived from [32], are specified in Section S17 in S1 File.

In the first scenario, we assume that all system variables are observable. The observations are generated by numerically integrating Eq. (15) from 0 to 6 min, using the parameters and initial conditions specified in Section S17 in S1 File. The trajectory is sampled every 0.8 min (21 observation time points). We consider observation datasets with and without noise, labeled as *DS*_0.00_ and *DS*_0.05_ respectively. The noise dataset (*DS*_0.05_) is generated as in the two previous test cases. We execute the pipeline, switching to stiff configurations (as described for the cell apoptosis case) after observing the first trials of the initial step. The tuned hyperparameters are listed in Table S16 in S1 File. The trained HNODE models fit the observations (Fig 9 for the dynamics of the variables *y*_1_ and *y*_2_, Figs S11 and S12 in S1 File for all the model variables).

**Fig 9.**
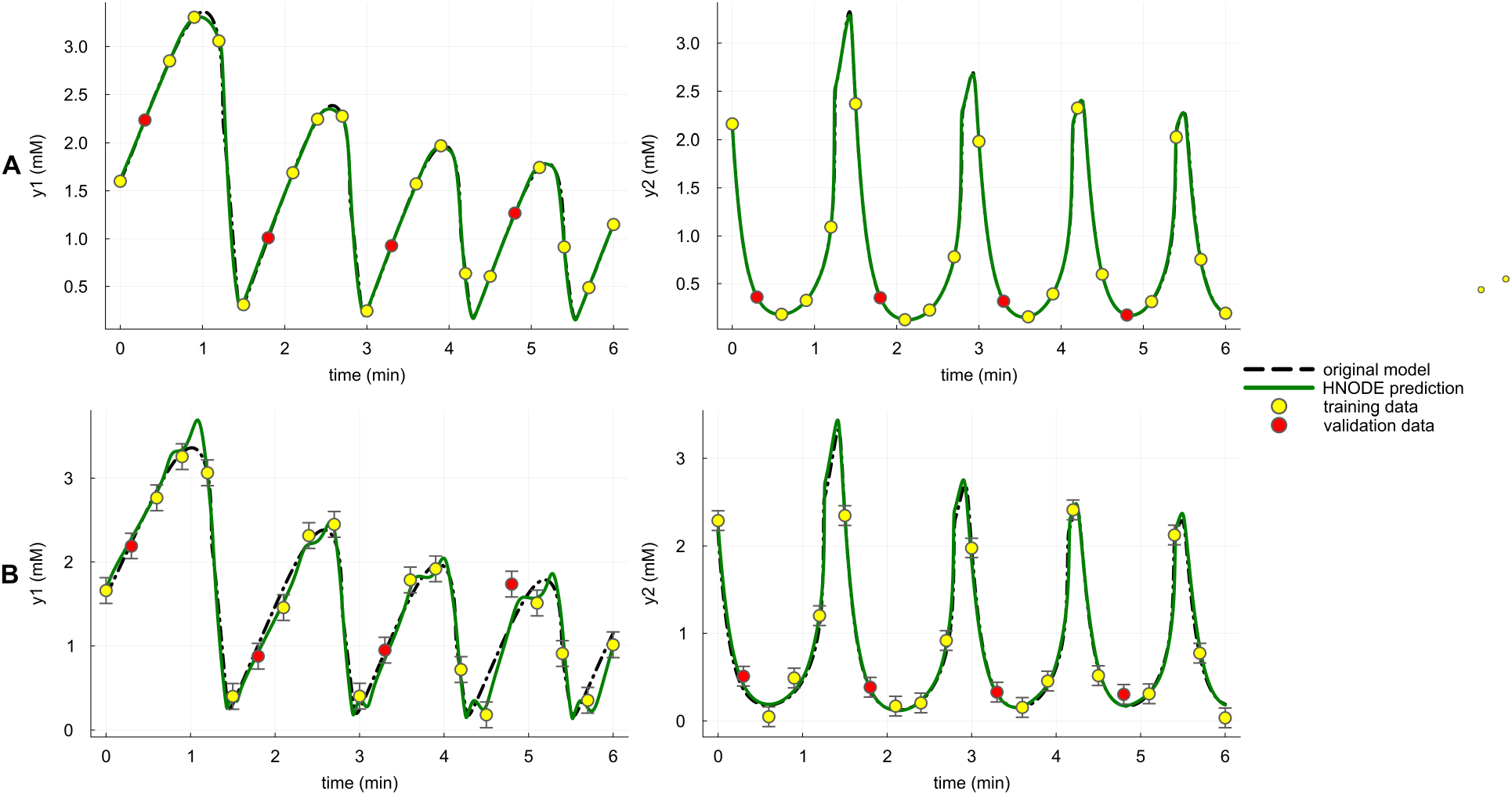
Dynamics predicted by the HNODE model in the yeast glycolysis test case, first scenario. The dynamics of *y*_1_ and *y*_2_ predicted by the HNODE model trained on *DS*_0.00_ and *DS*_0.05_ (shown in panels A and B respectively) are compared with the original model. The points represent the observations of the system, divided into training and validation sets.

In the noiseless case, the pipeline assesses the identifiability of all the parameters, whereas, in the presence of noise, *k*_1_ is classified as not identifiable. By comparing the norm of the projections of the mechanistic parameters onto the null space of *H*_*χ*_ (Fig S13 in S1 File), we can notice that also when estimated on *DS*_0.00_, the norm of the *k*_1_ projection falls just below our identifiability threshold. In both cases, the projections of *k*_1_ are composed by *k*_1_ itself and by the neural network parameters, thus embodying a potential compensation between *k*_1_ and the neural network. This is plausible since the parameter *k*_1_ in the HNODE model directly multiplies the variable *y*_1_ parameterized by the neural network.

The identifiable parameters are estimated with a maximum relative error of 1.94% (Table 5) on *DS*_0.00_, with the ground truth value of the parameter always falling within the estimated CIs. When estimated on *DS*_0.05_, the maximum relative error of the identifiable parameter estimates is 52.60%, with the CIs containing the relative ground truth value for 11 out of 12 parameters. Given the challenge of discerning whether the relative error of parameter estimates originates from the lack of mechanistic knowledge or from the noise in the observation dataset, we attempted to estimate all parameter values using the fully mechanistic model on dataset *DS*_0.05_. The outcomes indicate that the mean and maximum relative errors obtained with our pipeline are consistent with what is obtained with the fully mechanistic model (Section S23 in S1 File).

**Table 5.** Yeast glycolysis, first scenario: estimated values of mechanistic parameters.

| Par. | Target | Dataset | Estimate | Rel. err. | Id. | CI |
| --- | --- | --- | --- | --- | --- | --- |
| $k_1$ | $1.00 \cdot 10^2$ | $DS_{0.00}$ | $1.02 \cdot 10^2$ | 1.94% | ✓ | $(8.46 \cdot 10^1, 1.19 \cdot 10^2)$ |
| | | $DS_{0.05}$ | $1.25 \cdot 10^2$ | 25.43% | | |
| $K_1$ | $5.20 \cdot 10^{-1}$ | $DS_{0.00}$ | $5.25 \cdot 10^{-1}$ | 0.95% | ✓ | $(4.86 \cdot 10^{-1}, 5.67 \cdot 10^{-1})$ |
| | | $DS_{0.05}$ | $4.65 \cdot 10^{-1}$ | 10.61% | ✓ | $(1.58 \cdot 10^{-1}, 1.37)$ |
| $q$ | 4.00 | $DS_{0.00}$ | 4.04 | 1.02% | ✓ | (3.87, 4.21) |
| | | $DS_{0.05}$ | 3.79 | 5.33% | ✓ | (2.16, 5.42) |
| $k_2$ | 6.00 | $DS_{0.00}$ | 5.96 | 0.68% | ✓ | (4.68, 7.24) |
| | | $DS_{0.05}$ | 9.16 | 52.60% | ✓ | $(9.16 \cdot 10^{-1}, 1.99 \cdot 10^1)$ |
| $N$ | 1.00 | $DS_{0.00}$ | 1.00 | 0.13% | ✓ | $(8.20 \cdot 10^{-1}, 1.18)$ |
| | | $DS_{0.05}$ | $7.04 \cdot 10^{-1}$ | 29.61% | ✓ | $(1.0 \cdot 10^{-1}, 1.34)$ |
| $k_6$ | $1.20 \cdot 10^1$ | $DS_{0.00}$ | $1.19 \cdot 10^1$ | 0.57% | ✓ | $(1.16 \cdot 10^1, 1.23 \cdot 10^1)$ |
| | | $DS_{0.05}$ | 9.84 | 17.97% | ✓ | $(7.11, 1.26 \cdot 10^1)$ |
| $k_3$ | $1.60 \cdot 10^1$ | $DS_{0.00}$ | $1.59 \cdot 10^1$ | 0.59% | ✓ | $(1.56 \cdot 10^1, 1.62 \cdot 10^1)$ |
| | | $DS_{0.05}$ | $1.38 \cdot 10^1$ | 13.76% | ✓ | $(1.12 \cdot 10^1, 1.64 \cdot 10^1)$ |
| $A$ | 4.00 | $DS_{0.00}$ | 4.00 | 0.02% | ✓ | (3.98, 4.02) |
| | | $DS_{0.05}$ | 4.43 | 10.69% | ✓ | (4.21, 4.65) |
| $k_4$ | $1.00 \cdot 10^2$ | $DS_{0.00}$ | $9.94 \cdot 10^1$ | 0.64% | ✓ | $(9.73 \cdot 10^1, 1.01 \cdot 10^2)$ |
| | | $DS_{0.05}$ | $1.02 \cdot 10^2$ | 1.75% | ✓ | $(8.50 \cdot 10^1, 1.18 \cdot 10^2)$ |
| $\kappa$ | $1.30 \cdot 10^1$ | $DS_{0.00}$ | $1.29 \cdot 10^1$ | 0.49% | ✓ | $(1.26 \cdot 10^1, 1.33 \cdot 10^1)$ |
| | | $DS_{0.05}$ | $1.15 \cdot 10^1$ | 11.84% | ✓ | $(8.98, 1.39 \cdot 10^1)$ |
| $k_5$ | 1.28 | $DS_{0.00}$ | 1.28 | 0.03% | ✓ | (1.26, 1.30) |
| | | $DS_{0.05}$ | 1.31 | 2.40% | ✓ | (1.17, 1.45) |
| $\psi$ | $1.00 \cdot 10^{-1}$ | $DS_{0.00}$ | $1.00 \cdot 10^{-1}$ | 0.09% | ✓ | $(9.55 \cdot 10^{-2}, 1.05 \cdot 10^{-1})$ |
| | | $DS_{0.05}$ | $1.31 \cdot 10^{-1}$ | 30.55% | ✓ | $(8.13 \cdot 10^{-2}, 1.80 \cdot 10^{-1})$ |
| $k$ | 1.80 | $DS_{0.00}$ | 1.80 | 0.02% | ✓ | (1.76, 1.84) |
| | | $DS_{0.05}$ | 2.08 | 15.78% | ✓ | (1.68, 2.48) |

The symbol ✓ denotes the identifiable parameters according to our method. CIs for the model trained on $DS_{0.00}$ are computed assuming an uncertainty equal to 0.5% of the min-max of each time series.

In the second scenario, we assume that only the variables *y*_5_ and *y*_6_ are observable. We chose these variables because, in the original model, all parameters are identifiable when *y*_5_ and *y*_6_ are observed [7]. Following a consistent approach with the previous test case, the *in silico* observations are generated as in the first scenario, with the sampling frequency halved to 0.4 minutes, resulting in 42 observed time points. The pipeline is run analogously to the first scenario. The tuned hyperparameters are listed in Table S17 in S1 File, and the dynamics predicted by the HNODE model (Figs S14 and S15 in S1 File) fit the experimental data.

Similar to the first scenario, the results regarding parameter identifiability vary when the parameters are estimated on *DS*_0.00_ and *DS*_0.05_ (Table 6). When the parameters are estimated using the dataset without noise, the pipeline classifies six parameters as identifiable. However, when estimated using *DS*_0.05_, the parameters *K*_1_ and *k*_4_, classified as identifiable on *DS*_0.00_, are classified as non-identifiable. By analyzing the projections of the parameters onto the null subspace of *H*_*χ*_ (Fig 10), we can notice that *K*_1_ is classified as non-identifiable on *DS*_0.05_, but its projection falls just above the threshold. Conversely, parameter *k*_4_ is classified as identifiable on *DS*_0.00_, yet its projection falls just below the threshold. Interestingly, on both datasets, the neural network parameters significantly contribute to the projection of nearly all non-identifiable mechanistic parameters, except for *k*_2_ and *N* . This observation is consistent with our findings on the cell apoptosis test case: when the unknown variable *y*_1_ is not observable, the neural network can compensate for changes in mechanistic parameters, also not directly linked to *y*_1_ in the HNODE model.

**Table 6.** Yeast glycolysis, second scenario: estimated values of mechanistic parameters.

| Par. | Target | Dataset | Estimate | Rel. err. | Id. | CI |
| --- | --- | --- | --- | --- | --- | --- |
| $k_1$ | $1.00 \cdot 10^2$ | $DS_{0.00}$ | $5.42 \cdot 10^1$ | 45.77% | | |
| | | $DS_{0.05}$ | $7.02 \cdot 10^1$ | 29.84% | | |
| $K_1$ | $5.20 \cdot 10^{-1}$ | $DS_{0.00}$ | $5.20 \cdot 10^{-1}$ | 0.04% | ✓ | $(2.50 \cdot 10^{-1}, 1.08)$ |
| | | $DS_{0.05}$ | $7.91 \cdot 10^{-1}$ | 52.12% | | |
| $q$ | 4.00 | $DS_{0.00}$ | 3.63 | 9.26% | ✓ | (1.49, 5.76) |
| | | $DS_{0.05}$ | 4.44 | 11.05% | ✓ | $(9.45 \cdot 10^{-1}, 7.94)$ |
| $k_2$ | 6.00 | $DS_{0.00}$ | $1.04 \cdot 10^1$ | 74.00% | | |
| | | $DS_{0.05}$ | 2.52 | 57.96% | | |
| $N$ | 1.00 | $DS_{0.00}$ | $6.61 \cdot 10^{-1}$ | 33.94% | | |
| | | $DS_{0.05}$ | 2.26 | 126.02% | | |
| $k_6$ | $1.20 \cdot 10^1$ | $DS_{0.00}$ | $1.17 \cdot 10^1$ | 2.49% | ✓ | $(9.58, 1.38 \cdot 10^1)$ |
| | | $DS_{0.05}$ | $1.07 \cdot 10^1$ | 10.74% | ✓ | $(4.71, 1.67 \cdot 10^1)$ |
| $k_3$ | $1.60 \cdot 10^1$ | $DS_{0.00}$ | $1.63 \cdot 10^1$ | 1.61% | ✓ | $(1.29 \cdot 10^1, 1.96 \cdot 10^1)$ |
| | | $DS_{0.05}$ | $5.98 \cdot 10^1$ | 273.76% | | |
| $A$ | 4.00 | $DS_{0.00}$ | 4.00 | 0.00% | ✓ | (3.90, 4.10) |
| | | $DS_{0.05}$ | 3.79 | 5.27% | ✓ | (3.39, 4.19) |
| $k_4$ | $1.00 \cdot 10^2$ | $DS_{0.00}$ | $1.06 \cdot 10^2$ | 6.26% | | |
| | | $DS_{0.05}$ | $1.94 \cdot 10^2$ | 93.70% | | |
| $\kappa$ | $1.30 \cdot 10^1$ | $DS_{0.00}$ | $1.19 \cdot 10^1$ | 8.83% | | |
| | | $DS_{0.05}$ | $4.35 \cdot 10^1$ | 234.70% | | |
| $k_5$ | 1.28 | $DS_{0.00}$ | 1.33 | 3.74% | ✓ | (1.25, 1.41) |
| | | $DS_{0.05}$ | 1.35 | 5.48% | ✓ | (1.05, 1.65) |
| $\psi$ | $1.00 \cdot 10^{-1}$ | $DS_{0.00}$ | $1.17 \cdot 10^{-1}$ | 16.89% | | |
| | | $DS_{0.05}$ | $3.97 \cdot 10^{-1}$ | 297.04% | | |
| $k$ | 1.80 | $DS_{0.00}$ | 2.21 | 23.02% | | |
| | | $DS_{0.05}$ | 6.65 | 269.66% | | |

The symbol ✓ denotes the identifiable parameters according to our method. CIs for the model trained on $DS_{0.00}$ are computed assuming an uncertainty equal to 0.5% of the min-max of each time series.

**Fig 10.**
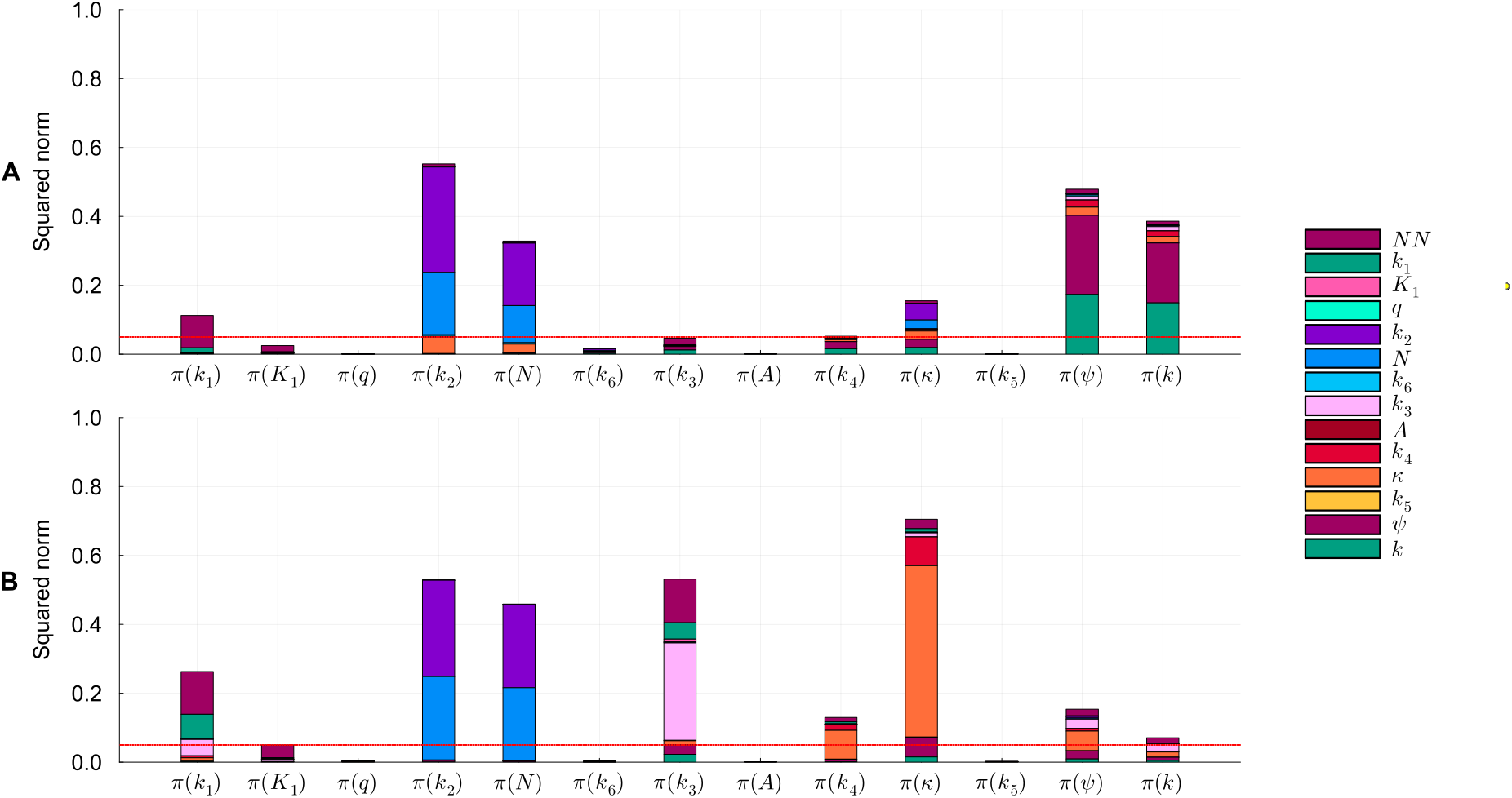
Identifiability analysis of mechanistic parameters in the yeast glycolysis test case, second scenario. Squared norm of the projections of the mechanistic parameters onto the null subspace of *H*_*χ*_ for the model trained on *DS*_0.00_ and *DS*_0.05_ (panel A and B respectively). The total height of the bar corresponds to the squared norm of the projection, while the different components of the projection are depicted in different colors. The red line indicates the threshold to determine the identifiability of the parameter (0.05).

The identifiable parameters are estimated with a maximum relative error of 9.26% on *DS*_0.00_. On the dataset with error *DS*_0.05_ the identifiable parameters are estimated with a maximum relative error of 11.05%. In both cases, the ground truth value consistently falls within the confidence interval estimated.

## Discussion

We have introduced a pipeline for estimating mechanistic parameters and discussing their local identifiability within an incompletely specified mechanistic model. This workflow leverages the HNODE framework to embed the original model into a larger model, in which the unknown portions of the system are described by neural networks. The primary novelties of our work lie in our approach to conducting a global exploration of the mechanistic parameter search space and our method to evaluate the parameter identifiability. First, we include the initial values of mechanistic parameters into the hyperparameters to be tuned. This involves combining Bayesian Optimization with gradient-based search, with the goal of globally exploring the parameter search space. Second, to assess the local identifiability at-a-point of the parameters, we extend a classical approach for identifiability analysis in mechanistic models. Notably, identifiability analysis for HNODE models has not been previously investigated in the literature, which has predominantly focused on ensuring the identifiability of the mechanistic component through regularization of the training cost function [28, 30].

The primary limitations of our work include: (1) the assumption of having access to initial conditions for all state variables within systems, even under partial observability; (2) the computational cost associated with the hyperparameter tuning phase; (3) the intrinsic local nature of identifiability analysis; (4) the arbitrary selection of hyperparameters, *ϵ* and *δ*, for identifiability analysis; and (5) the methodology employed for estimating confidence intervals. All these limitations are discussed in the following paragraphs.

We assume knowledge of the initial states of all system variables, possibly with noise, even when only a subset of them is observable. In real-world scenarios, this assumption may not always be valid. While in certain cases, scientists have concrete physical insights into the system’s initial state, in others, initial concentrations need to be estimated alongside the parameters. Notably, this limitation is shared by other approaches to system identification [7].

The initial phase of hyperparameter tuning and exploration of the mechanistic search space is the most computationally expensive step within the pipeline. We employed the TPE algorithm, and this algorithm has proven effective even in high-dimensional parameter search spaces, as evidenced by the Yeast Glycolysis test case, featuring 13 mechanistic parameters and 6 hyperparameters. However, employing the TPE algorithm entails sequentially repeating the training of the candidate HNODE model, each time with a limited number of epochs. It is important to note that the problem of hyperparameter tuning is a long-standing problem and different approaches have been proposed for this goal [59]. These approaches range from random and grid search to Bayesian optimization and genetic algorithms: in this work, we did not conduct a comparative analysis of TPE performance against other hyperparameter tuning methods.

The local nature of our identifiability analysis implies that it might overlook parameter non-identifiability if distinct parameterizations leading to similar model simulations are isolated in the search space, as separate local minima of the cost function.

The hyperparameters *ϵ* and *δ* used for identifiability analysis, although related, have different meanings. The threshold *ϵ* intuitively distinguishes what we assume to be negligible changes in the HNODE model behavior from non-negligible changes. This hyperparameter is also present in existing Hessian-based methods for identifiability analysis of mechanistic models [44]. The threshold *δ*, introduced in our method, intuitively quantifies how much it is possible to perturb the mechanistic parameter with a negligible effect on the model simulations. *δ* has been introduced to overcome the limitation of the dominant parameter approach in the context of UDE. Existing Hessian-based methods for identifiability analysis of mechanistic models [44] primarily categorize parameters as non-identifiable if they have the highest projection (in absolute value) onto eigenvectors associated with null eigenvalues. We decided against using this approach because we observed empirically that in HNODE models when compensation occurs between the neural network and mechanistic parameters, the dominant parameter in the null direction is typically a neural network parameter. Thus, considering only the dominant parameter, we risk overlooking the role of the mechanistic parameter. In our test cases, we employed *ϵ* = 10^−5^ and *δ* = 0.05. The results of the first two test cases are quite robust to the choice of *ϵ* and *δ* (Sections S4 and S16 in S1 File). However, the third test case is more complex: here changes in *ϵ* and *δ* would lead to different results for certain parameters (Section S24 in S1 File).

The fifth limitation of our work is the CI estimation. The FIM-based approach we employed has several limitations [60]. By using it, we implicitly assume the unbiasedness and normality of our estimators [49]. Additionally, since our estimators are not linear, the CIs estimated are only lower bounds of the real CIs [49]. We opted for this method because of its computational advantages over other methods, such as likelihood profiling or bootstrapping-based methods [15, 60], which would necessitate multiple trainings of the HNODE model.

We tested our pipeline in various *in silico* scenarios of increasing complexity, assuming a partial lack of mechanistic knowledge in three models acknowledged as benchmarks in computational biology. These tests encompass various conditions, including different levels of noise in the training data and different assumptions regarding the observability of the system variables. Across all the examined scenarios, our pipeline consistently enabled the analysis of parameter identifiability and the accurate estimation of identifiable parameters.

Despite its limitations, the proposed workflow represents an initial step toward adapting traditional methods utilized in entirely mechanistic models to the HNODE modeling scenario. In future works, we aim to address the limitations of the pipeline. Firstly, we will consider different state-of-the-art approaches for hyperparameter tuning, such as Gaussian process [61], evolutionary [59], and genetic approaches [62], comparing their performances with TPE. Secondly, we plan to expand the number of test cases to derive a less arbitrary choice of the thresholds *ϵ* and *δ*, which could vary based on the number of parameters in the HNODE model. Thirdly, we intend to compare the FIM-based method for estimating CIs with other approaches that have demonstrated greater reliability in completely mechanistic models, as described in [15, 60].

## Supporting information

**S1 File Supplementary sections, tables and figures**.

## References

1. Motta S, Pappalardo F. Mathematical modeling of biological systems. Briefings in Bioinformatics. 2013;14(4):411–422.

2. Mogilner A, Wollman R, Marshall WF. Quantitative modeling in cell biology: what is it good for? Developmental cell. 2006;11(3):279–287.

3. Gábor A, Banga JR. Robust and efficient parameter estimation in dynamic models of biological systems. BMC systems biology. 2015;9(1):1–25.

4. Fischer HP. Mathematical modeling of complex biological systems: from parts lists to understanding systems behavior. Alcohol Research & Health. 2008;31(1):49.

5. Baker RE, Pena JM, Jayamohan J, Jérusalem A. Mechanistic models versus machine learning, a fight worth fighting for the biological community? Biology letters. 2018;14(5):20170660.

6. Cornish-Bowden A. Fundamentals of enzyme kinetics. John Wiley & Sons; 2013.

7. Yazdani A, Lu L, Raissi M, Karniadakis GE. Systems biology informed deep learning for inferring parameters and hidden dynamics. PLoS computational biology. 2020;16(11):e1007575.

8. Lillacci G, Khammash M. Parameter estimation and model selection in computational biology. PLoS computational biology. 2010;6(3):e1000696.

9. Sun J, Garibaldi JM, Hodgman C. Parameter estimation using metaheuristics in systems biology: a comprehensive review. IEEE/ACM transactions on computational biology and bioinformatics. 2011;9(1):185–202.

10. Mendes P, Kell D. Non-linear optimization of biochemical pathways: applications to metabolic engineering and parameter estimation. Bioinformatics (Oxford, England). 1998;14(10):869–883.

11. Reali F, Priami C, Marchetti L. Optimization algorithms for computational systems biology. Frontiers in Applied Mathematics and Statistics. 2017;3:6.

12. Liepe J, Kirk P, Filippi S, Toni T, Barnes CP, Stumpf MP. A framework for parameter estimation and model selection from experimental data in systems biology using approximate Bayesian computation. Nature protocols. 2014;9(2):439–456.

13. Linden NJ, Kramer B, Rangamani P. Bayesian parameter estimation for dynamical models in systems biology. PLOS Computational Biology. 2022;18(10):e1010651.

14. Meskin N, Nounou H, Nounou M, Datta A, Dougherty ER. Parameter estimation of biological phenomena modeled by S-systems: an extended Kalman filter approach. In: 2011 50th IEEE Conference on Decision and Control and European Control Conference. IEEE; 2011. p. 4424–4429.

15. Kreutz C, Raue A, Kaschek D, Timmer J. Profile likelihood in systems biology. The FEBS journal. 2013;280(11):2564–2571.

16. Anstett-Collin F, Denis-Vidal L, Millérioux G. A priori identifiability: An overview on definitions and approaches. Annual Reviews in Control. 2020;50:139–149.

17. Lam NN, Docherty PD, Murray R. Practical identifiability of parametrised models: A review of benefits and limitations of various approaches. Mathematics and Computers in Simulation. 2022;199:202–216.

18. Yeo HC, Selvarajoo K. Machine learning alternative to systems biology should not solely depend on data. Briefings in Bioinformatics. 2022;23(6):bbac436.

19. Engelhardt B, Frőhlich H, Kschischo M. Learning (from) the errors of a systems biology model. Scientific reports. 2016;6(1):20772.

20. Zou BJ, Levine ME, Zaharieva DP, Johari R, Fox EB. Hybrid Square Neural ODE Causal Modeling. arXiv preprint arXiv:240217233. 2024;.

21. Lanzieri D, Lanusse F, Starck JL. Hybrid Physical-Neural ODEs for Fast N-body Simulations. arXiv preprint arXiv:220705509. 2022;.

22. Grigorian G, George SV, Lishak S, Shipley RJ, Arridge S. A hybrid neural ordinary differential equation model of the cardiovascular system. Journal of the Royal Society Interface. 2024;21(212):20230710.

23. Alber M, Buganza Tepole A, Cannon WR D. S, Dura-Bernal S, Garikipati K, et al. Integrating machine learning and multiscale modeling—perspectives, challenges, and opportunities in the biological, biomedical, and behavioral sciences. NPJ digital medicine. 2019;2(1):115.

24. Zhang T, Androulakis IP, Bonate P, Cheng L, Helikar T, Parikh J, et al. Two heads are better than one: current landscape of integrating QSP and machine learning: an ISoP QSP SIG white paper by the working group on the integration of quantitative systems pharmacology and machine learning. Journal of Pharmacokinetics and Pharmacodynamics. 2022;49(1):5–18.

25. Rackauckas C, Ma Y, Martensen J, Warner C, Zubov K, Supekar R, et al. Universal differential equations for scientific machine learning. arXiv preprint arXiv:200104385. 2020;.

26. Bräm DS, Nahum U, Schropp J, Pfister M, Koch G. Low-dimensional neural ODEs and their application in pharmacokinetics. Journal of Pharmacokinetics and Pharmacodynamics. 2023; p. 1–18.

27. Valderrama D, Ponce-Bobadilla AV, Mensing S, Fröhlich H, Stodtmann S. Integrating machine learning with pharmacokinetic models: Benefits of scientific machine learning in adding neural networks components to existing PK models. CPT: Pharmacometrics & Systems Pharmacology. 2023;.

28. Takeishi N, Kalousis A. Deep Grey-Box Modeling With Adaptive Data-Driven Models Toward Trustworthy Estimation of Theory-Driven Models. In: International Conference on Artificial Intelligence and Statistics. PMLR; 2023. p. 4089–4100.

29. Kidger P. On neural differential equations. arXiv preprint arXiv:220202435. 2022;.

30. Yin Y, Le Guen V, Dona J, de Bézenac E, Ayed I, Thome N, et al. Augmenting physical models with deep networks for complex dynamics forecasting. Journal of Statistical Mechanics: Theory and Experiment. 2021;2021(12):124012.

31. Aldridge BB, Haller G, Sorger PK, Lauffenburger DA. Direct Lyapunov exponent analysis enables parametric study of transient signalling governing cell behaviour. IEE Proceedings-Systems Biology. 2006;153(6):425–432.

32. Ruoff P, Christensen MK, Wolf J, Heinrich R. Temperature dependency and temperature compensation in a model of yeast glycolytic oscillations. Biophysical chemistry. 2003;106(2):179–192.

33. Chen RT, Rubanova Y, Bettencourt J, Duvenaud DK. Neural ordinary differential equations. Advances in neural information processing systems. 2018;31.

34. Errico RM. What is an adjoint model? Bulletin of the American Meteorological Society. 1997;78(11):2577–2592.

35. Allaire G. A review of adjoint methods for sensitivity analysis, uncertainty quantification and optimization in numerical codes. Ingénieurs de l’Automobile. 2015;836:33–36.

36. Ghosh A, Behl H, Dupont E, Torr P, Namboodiri V. Steer: Simple temporal regularization for neural ode. Advances in Neural Information Processing Systems. 2020;33:14831–14843.

37. Kim S, Ji W, Deng S, Ma Y, Rackauckas C. Stiff neural ordinary differential equations. Chaos: An Interdisciplinary Journal of Nonlinear Science. 2021;31(9):093122.

38. Turan EM, Jäschke J. Multiple shooting for training neural differential equations on time series. IEEE Control Systems Letters. 2021;6:1897–1902.

39. Kingma DP, Ba J. Adam: A method for stochastic optimization. arXiv preprint arXiv:14126980. 2014;.

40. Liu DC, Nocedal J. On the limited memory BFGS method for large scale optimization. Mathematical programming. 1989;45(1-3):503–528.

41. Gao Y, Yu T, Li J. Bayesian optimization with local search. In: Machine Learning, Optimization, and Data Science: 6th International Conference, LOD 2020, Siena, Italy, July 19–23, 2020, Revised Selected Papers, Part II 6. Springer; 2020. p. 350–361.

42. Bergstra J, Bardenet R, Bengio Y, Kégl B. Algorithms for hyper-parameter optimization. Advances in neural information processing systems. 2011;24.

43. Falkner S, Klein A, Hutter F. BOHB: Robust and efficient hyperparameter optimization at scale. In: International conference on machine learning. PMLR; 2018. p. 1437–1446.

44. Quaiser T, Mönnigmann M. Systematic identifiability testing for unambiguous mechanistic modeling–application to JAK-STAT, MAP kinase, and NF-κ B signaling pathway models. BMC systems biology. 2009;3(1):1–21.

45. Gutenkunst RN, Waterfall JJ, Casey FP, Brown KS, Myers CR, Sethna JP. Universally sloppy parameter sensitivities in systems biology models. PLoS computational biology. 2007;3(10):e189.

46. Transtrum MK, Machta BB, Brown KS, Daniels BC, Myers CR, Sethna JP. Perspective: Sloppiness and emergent theories in physics, biology, and beyond. The Journal of chemical physics. 2015;143(1).

47. Jagadeesan P, Raman K, Tangirala AK. Sloppiness: Fundamental study, new formalism and its application in model assessment. Plos one. 2023;18(3):e0282609.

48. Rodriguez-Fernandez M, Banga JR, Doyle III FJ. Novel global sensitivity analysis methodology accounting for the crucial role of the distribution of input parameters: application to systems biology models. International Journal of Robust and Nonlinear Control. 2012;22(10):1082–1102.

49. Tangirala AK. Principles of system identification: theory and practice. Crc Press; 2018.

50. Stoica P, Marzetta TL. Parameter estimation problems with singular information matrices. IEEE Transactions on Signal Processing. 2001;49(1):87–90.

51. Glorot X, Bengio Y. Understanding the difficulty of training deep feedforward neural networks. In: Proceedings of the thirteenth international conference on artificial intelligence and statistics. JMLR Workshop and Conference Proceedings; 2010. p. 249–256.

52. Bezanson J, Edelman A, Karpinski S, Shah VB. Julia: A fresh approach to numerical computing. SIAM review. 2017;59(1):65–98.

53. Akiba T, Sano S, Yanase T, Ohta T, Koyama M. Optuna: A next-generation hyperparameter optimization framework. In: Proceedings of the 25th ACM SIGKDD international conference on knowledge discovery & data mining; 2019. p. 2623–2631.

54. Wangersky PJ. Lotka-Volterra population models. Annual Review of Ecology and Systematics. 1978;9(1):189–218.

55. Verner JH. Numerically optimal Runge–Kutta pairs with interpolants. Numerical Algorithms. 2010;53(2-3):383–396.

56. Hosea M, Shampine L. Analysis and implementation of TR-BDF2. Applied Numerical Mathematics. 1996;20(1-2):21–37.

57. Schmidt M, Lipson H. Distilling free-form natural laws from experimental data. science. 2009;324(5923):81–85.

58. Brunton SL, Proctor JL, Kutz JN. Discovering governing equations from data by sparse identification of nonlinear dynamical systems. Proceedings of the national academy of sciences. 2016;113(15):3932–3937.

59. Vincent AM, Jidesh P. An improved hyperparameter optimization framework for AutoML systems using evolutionary algorithms. Scientific Reports. 2023;13(1):4737.

60. Joshi M, Seidel-Morgenstern A, Kremling A. Exploiting the bootstrap method for quantifying parameter confidence intervals in dynamical systems. Metabolic engineering. 2006;8(5):447–455.

61. Wu J, Chen XY, Zhang H, Xiong LD, Lei H, Deng SH. Hyperparameter optimization for machine learning models based on Bayesian optimization. Journal of Electronic Science and Technology. 2019;17(1):26–40.

62. Aszemi NM, Dominic P. Hyperparameter optimization in convolutional neural network using genetic algorithms. International Journal of Advanced Computer Science and Applications. 2019;10(6).

