## Supplementary File S1 for "Robust parameter estimation and identifiability analysis with Hybrid Neural Ordinary Differential Equations in Computational Biology"

### S1 file: Supplementary sections, tables and figures

#### S1 Lotka-Volterra test case: experimental setting

To simulate in silico observations for the Lotka-Volterra test case, the ODE system has been solved numerically employing the *Vern7* solver from the *DifferentialEquations.jl* library. The solver settings are configured with an absolute tolerance of $10^{-7}$ and a relative tolerance of $10^{-6}$. The parameters and initial states, derived from [RMM$^+$20], are detailed in Table S1 and S2.

Table S1: **Lotka-Volterra test case: parameters employed for generating the *in silico* dataset**

| **Parameter** | **Value** | **Unit** |
|---|---|---|
| $\alpha$ | 1.3 | year$^{-1}$ |
| $\beta$ | 0.9 | year$^{-1}$ |
| $\gamma$ | 0.8 | year$^{-1}$ |
| $\delta$ | 1.8 | year$^{-1}$ |

Table S2: **Lotka-Volterra - initial states employed for generating the *in silico* dataset**

| **Variable** | **Value** |
|---|---|
| $y_1$ | 3.416 |
| $y_2$ | 1.537 |

### S2 Lotka-Volterra test case: tuned hyper-parameters

Table S3: **Lotka-Volterra test case: tuned hyperparameters of the HNODE model.**

| Name | Search Space | Tuned value $DS_{0.00}$ | Tuned value $DS_{0.05}$ |
|---|---|---|---|
| *Neural network architecture* | | | |
| hidden layer number | $\{2, 3\}$ | 2 | 3 |
| hidden layer width | $\{4, 8, 16, 32\}$ | 16 | 8 |
| *Starting values for mechanistic parameters* | | | |
| $\alpha$ | $[1.30 \cdot 10^{-2}, 1.30 \cdot 10^{2}]$ | $9.85 \cdot 10^{-1}$ | $5.81 \cdot 10^{-1}$ |
| *Training hyper-parameters* | | | |
| learning rate | $[10^{-5}, 10^{-1}]$ | $3.12 \cdot 10^{-2}$ | $7.10 \cdot 10^{-3}$ |
| $\lambda$ | $\{10^{-3}, 10^{-2}, 10^{-1}, 1\}$ | ND | 1.0 |
| $k$ (segmentation) | $\{2, 3, 4, 5, 6, 7, 8, 9, 10\}$ | 2 | 5 |
| $\rho$ | $[10^{-3}, 10^{3}]$ | $2.61 \cdot 10^{-3}$ | 6.52 |

### S3 Lotka-Volterra test case: attempt to retrieve the identifiability of $\alpha$ by regularizing the cost function

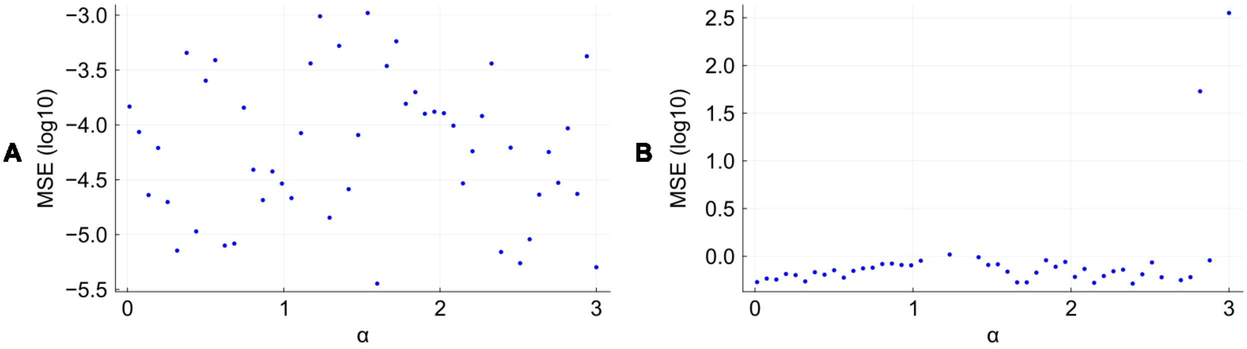


Figure S1: **Mean Squared Error of HNODE models trained with fixed $\alpha$ in the Lotka-Volterra test case.** Mean Squared Errors (MSE) of the HNODE model trained with varying fixed $\alpha$ on $DS_{0.00}$ and $DS_{0.05}$ (A and B respectively). On the x-axis, values of $\alpha$ are sampled in the interval $[0.013, 3.0]$. On the y-axis, the MSE of the dynamics predicted by the HNODE models with fixed $\alpha$ with respect to the observed data is depicted. In dataset $DS_{0.05}$, the MSE has been computed normalizing for the uncertainty. The correlation between $\alpha$ and $MSE$ is very weak in $DS_{0.00}$ (-0.037) and weak in $DS_{0.05}$ (0.293).

### S4 Lotka-Volterra test case: impact of $\epsilon$ and $\delta$ on the identifiability analysis results

To assess the impact of varying hyperparameters $\epsilon$ and $\delta$ on the identifiability analysis, we conducted the analysis using different values for these parameters (Table S4). Our findings indicate that the results exhibit robustness against changes in $\epsilon$ and $\delta$.

Table S4: **Lotka-Volterra test case: parameter identifiability with different choices of $\epsilon$ and $\delta$.**

| | | | $\epsilon = 10^{-6}$ | | | $\epsilon = 10^{-5}$ | | $\epsilon = 10^{-4}$ | |
|---|---|---|---|---|---|---|---|---|---|
| **Par.** | **Dataset** | $\delta = 0.03$ | $\delta = 0.05$ | $\delta = 0.07$ | $\delta = 0.03$ | $\delta = 0.05$ | $\delta = 0.07$ | $\delta = 0.05$ | $\delta = 0.07$ |
| $\alpha$ | $DS_{0.00}$ | | | | | | | | |
| | $DS_{0.05}$ | | | | | | | | |

The symbol ✓ denotes the parameters classified as identifiable with the $\epsilon$ and $\delta$ threshold specified in the header.

#### S5 Cell apoptosis test case: experimental setting

To simulate in silico observations, the cell apoptosis ODE system is solved numerically employing the *TRBDF2* solver from the *DifferentialEquations.jl* library. The solver settings are configured with an absolute tolerance of $10^{-7}$ and a relative tolerance of $10^{-6}$. The parameters and initial states, derived from [AHSL06], are detailed in Table S5 and S6.

Table S5: **Cell apoptosis test case: parameters employed for generating the *in silico* dataset**

| Parameter | Value | Unit |
|---|---|---|
| $k_1$ | 0.9612 | $\text{cell} \cdot (\text{h} \cdot 10^5 \cdot \text{molecules})^{-1}$ |
| $k_{d1}$ | 36.0 | $\text{h}^{-1}$ |
| $k_{d2}$ | 28.8 | $\text{h}^{-1}$ |
| $k_3$ | 24.48 | $\text{cell} \cdot (\text{h} \cdot 10^5 \cdot \text{molecules})^{-1}$ |
| $k_{d3}$ | 180.0 | $\text{h}^{-1}$ |
| $k_{d4}$ | 3.6 | $\text{h}^{-1}$ |
| $k_5$ | 25200.0 | $\text{cell} \cdot (\text{h} \cdot 10^5 \cdot \text{molecules})^{-1}$ |
| $k_{d5}$ | 0.06012 | $\text{h}^{-1}$ |
| $k_{d6}$ | 0.6012 | $\text{h}^{-1}$ |

Table S6: **Cell apoptosis test case: initial states employed for generating the *in silico* dataset**

| Variable | Value | Unit |
|---|---|---|
| $y_1$ | 1.34 | $10^5 \cdot \text{molecules} \cdot \text{cell}^{-1}$ |
| $y_2$ | 1.0 | $10^5 \cdot \text{molecules} \cdot \text{cell}^{-1}$ |
| $y_3$ | 2.67 | $10^5 \cdot \text{molecules} \cdot \text{cell}^{-1}$ |
| $y_4$ | 0.0 | $10^5 \cdot \text{molecules} \cdot \text{cell}^{-1}$ |
| $y_5$ | 0.0 | $10^5 \cdot \text{molecules} \cdot \text{cell}^{-1}$ |
| $y_6$ | 0.0 | $10^5 \cdot \text{molecules} \cdot \text{cell}^{-1}$ |
| $y_7$ | 0.029 | $10^5 \cdot \text{molecules} \cdot \text{cell}^{-1}$ |
| $y_8$ | 0.0 | $10^5 \cdot \text{molecules} \cdot \text{cell}^{-1}$ |

### S6 Cell apoptosis test case (first scenario): tuned hyper-parameters

Table S7: **Cell apoptosis test case (first scenario): tuned hyperparameters of the HNODE model.**

| Name | Search Space | Tuned value $DS_{0.00}$ | Tuned value $DS_{0.05}$ |
|---|---|---|---|
| *Neural network architecture* | | | |
| hidden layer number | $\{2, 3, 4, 5, 6\}$ | 2 | 2 |
| hidden layer width | $\{4, 8, 16, 32\}$ | 8 | 8 |
| *Starting values for mechanistic parameters* | | | |
| $k_1$ | $[9.61 \cdot 10^{-3}, 9.61 \cdot 10^{1}]$ | $3.30 \cdot 10^{1}$ | $7.30 \cdot 10^{1}$ |
| $k_{d1}$ | $[3.60 \cdot 10^{-1}, 3.60 \cdot 10^{3}]$ | $1.12 \cdot 10^{3}$ | $2.87 \cdot 10^{3}$ |
| $k_{d2}$ | $[2.88 \cdot 10^{-1}, 2.88 \cdot 10^{3}]$ | $4.77 \cdot 10^{1}$ | $1.51 \cdot 10^{3}$ |
| $k_3$ | $[2.45 \cdot 10^{-1}, 2.45 \cdot 10^{3}]$ | $1.67 \cdot 10^{2}$ | $2.05 \cdot 10^{3}$ |
| $k_{d3}$ | $[1.80, 1.80 \cdot 10^{4}]$ | $2.51 \cdot 10^{3}$ | $9.08 \cdot 10^{3}$ |
| $k_{d4}$ | $[3.60 \cdot 10^{-2}, 3.60 \cdot 10^{2}]$ | $2.06 \cdot 10^{2}$ | $8.41 \cdot 10^{1}$ |
| $k_5$ | $[2.52 \cdot 10^{2}, 2.52 \cdot 10^{6}]$ | $2.37 \cdot 10^{6}$ | $3.92 \cdot 10^{4}$ |
| $k_{d5}$ | $[6.01 \cdot 10^{-4}, 6.01]$ | $6.30 \cdot 10^{-1}$ | 3.58 |
| $k_{d6}$ | $[6.01 \cdot 10^{-3}, 6.01 \cdot 10^{1}]$ | $4.41 \cdot 10^{1}$ | $1.73 \cdot 10^{1}$ |
| *Training hyper-parameters* | | | |
| learning rate | $[10^{-5}, 10^{-1}]$ | $3.82 \cdot 10^{-2}$ | $2.01 \cdot 10^{-2}$ |
| $\lambda$ | $\{10^{-3}, 10^{-2}, 10^{-1}, 1\}$ | ND | 0.01 |
| $k$ (segmentation) | $\{2, 3, 4, 5, 6, 7, 8, 9, 10\}$ | 2 | 8 |
| $\rho$ | $[10^{-3}, 10^{3}]$ | $1.18 \cdot 10^{-3}$ | $1.11 \cdot 10^{1}$ |

### S7 Cell apoptosis test case (first scenario): identifiability analysis

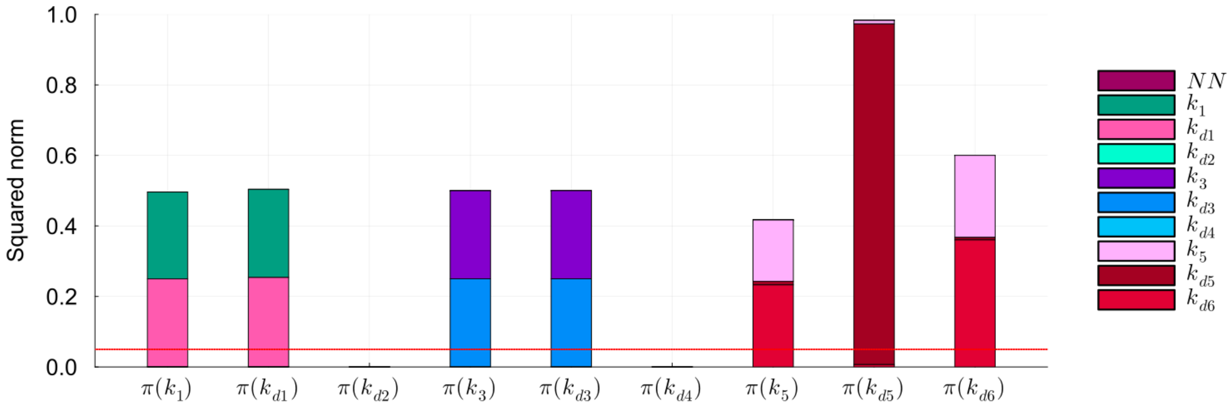


Figure S2: **Identifiability analysis of mechanistic parameters in the cell apoptosis test case, first scenario ($DS_{0.05}$).** Squared norm of the projections of the mechanistic parameters onto the null subspace of $H_\chi$ for the model trained on $DS_{0.05}$. The total height of the bar corresponds to the squared norm of the projection, while the different components of the projection are depicted in different colors. The red line indicates the threshold to determine the identifiability of the parameter (0.05).

#### S8 Cell apoptosis test case (first scenario): predicted dynamics

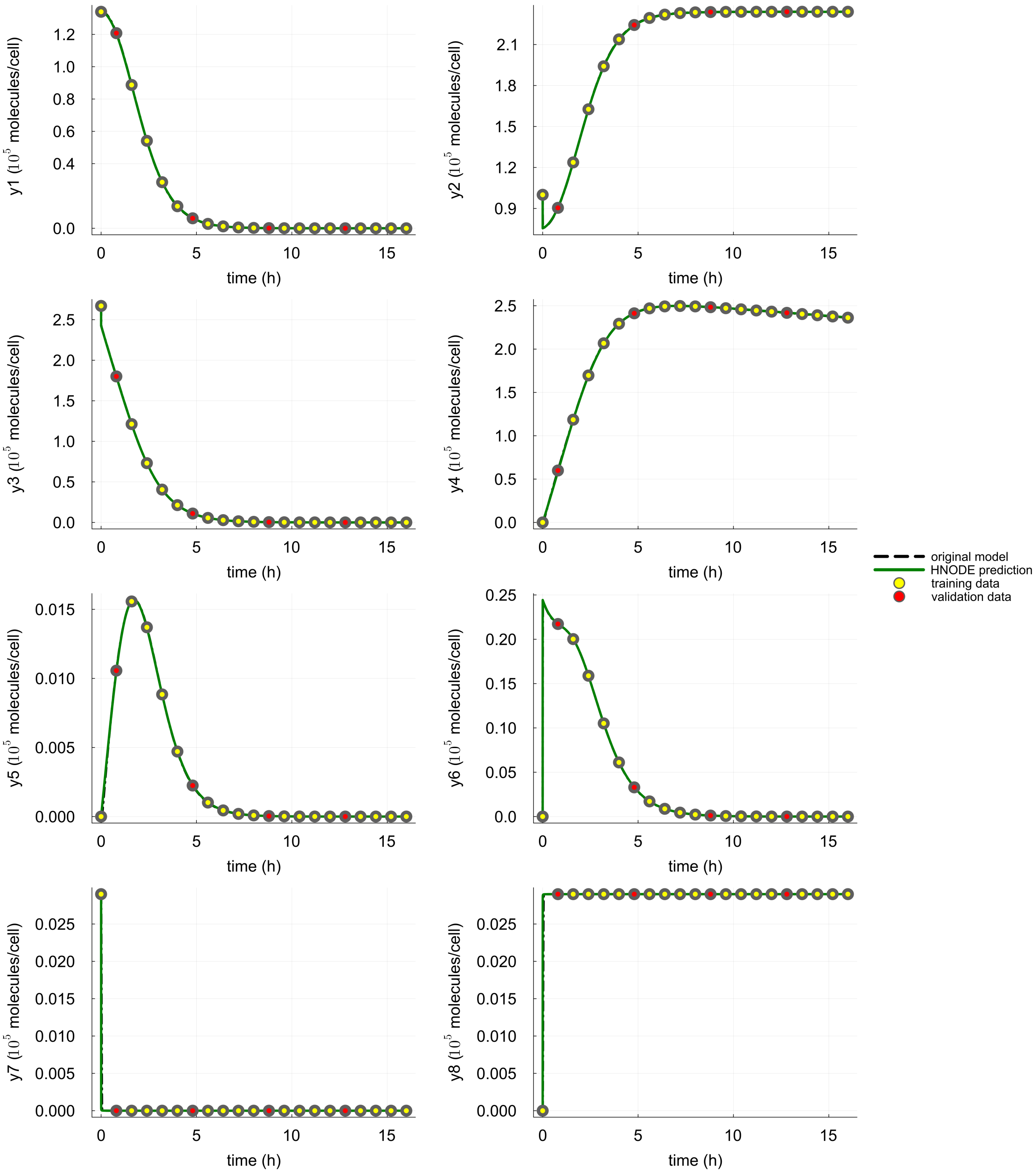


Figure S3: **Dynamics predicted by the HNODE model in the cell apoptosis test case, first scenario ($DS_{0.00}$).** The dynamics predicted by the HNODE model trained on $DS_{0.00}$ are compared with the original model. The points represent the observations of the system, divided into training and validation sets.

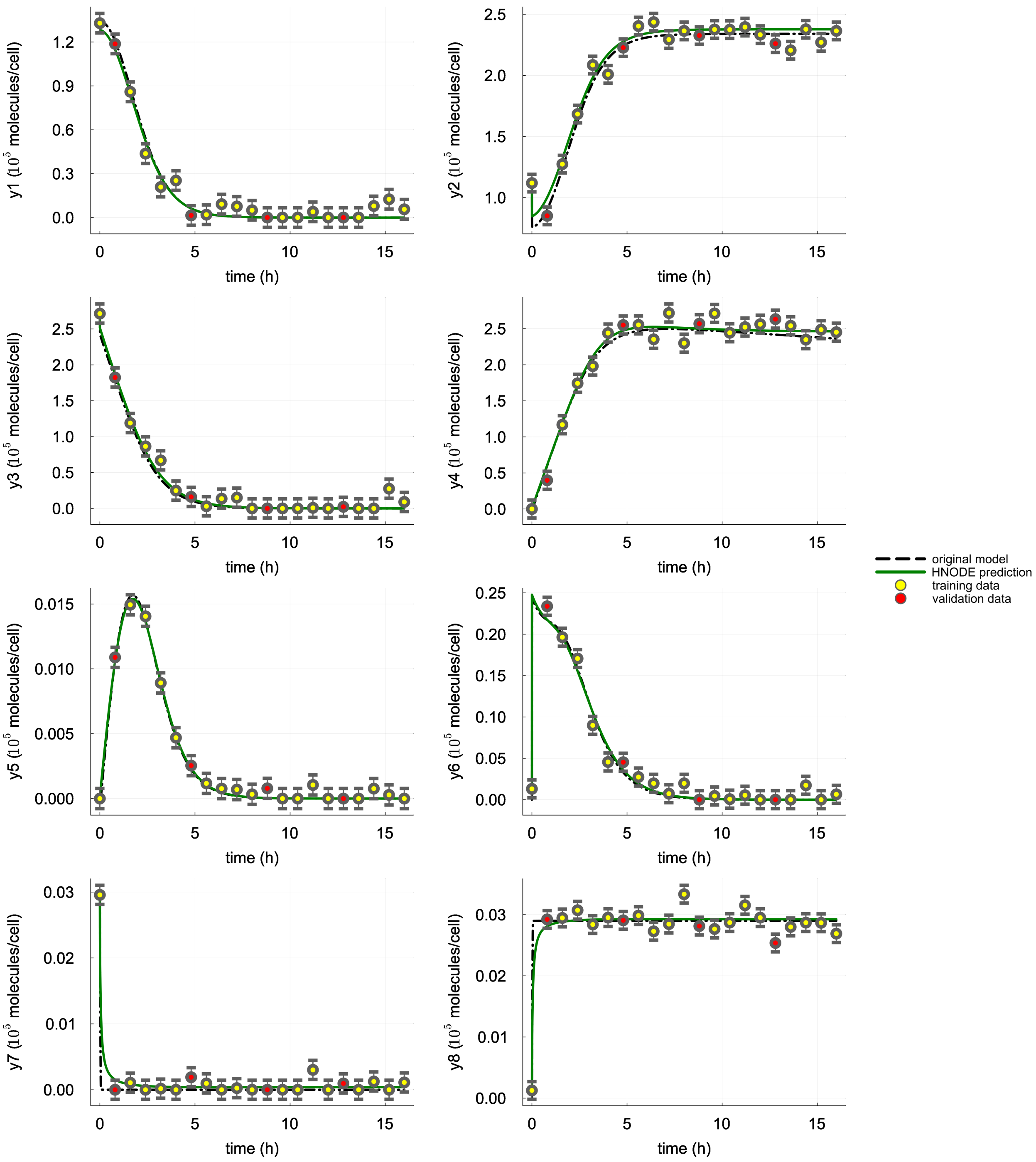


Figure S4: **Dynamics predicted by the HNODE model in the cell apoptosis test case, first scenario ($DS_{0.05}$).** The dynamics predicted by the HNODE model trained on $DS_{0.05}$ are compared with the original model. The points represent the observations of the system, divided into training and validation sets.

### S9 Cell apoptosis test case (first scenario, with $k_{d1}$ and $k_3$ fixed): tuned hyper-parameters

Table S8: **Cell apoptosis test case (first scenario, with $k_{d1}$ and $k_3$ fixed): tuned hyperparameters of the HNODE model.**

| **Name** | **Search Space** | **Tuned value** $DS_{0.00}$ | **Tuned value** $DS_{0.05}$ |
|---|---|---|---|
| *Neural network architecture* | | | |
| hidden layer number | $\{2, 3, 4, 5, 6\}$ | 2 | 4 |
| hidden layer width | $\{4, 8, 16, 32\}$ | 8 | 8 |
| *Starting values for mechanistic parameters* | | | |
| $k_1$ | $[9.61 \cdot 10^{-3}, 9.61 \cdot 10^{1}]$ | $1.53 \cdot 10^{1}$ | 8.58 |
| $k_{d2}$ | $[2.88 \cdot 10^{-1}, 2.88 \cdot 10^{3}]$ | $3.43 \cdot 10^{1}$ | $5.70 \cdot 10^{2}$ |
| $k_{d3}$ | $[1.80, 1.80 \cdot 10^{4}]$ | $2.74 \cdot 10^{3}$ | $1.28 \cdot 10^{4}$ |
| $k_{d4}$ | $[3.60 \cdot 10^{-2}, 3.60 \cdot 10^{2}]$ | $9.04 \cdot 10^{1}$ | $8.42 \cdot 10^{1}$ |
| $k_5$ | $[2.52 \cdot 10^{2}, 2.52 \cdot 10^{6}]$ | $2.05 \cdot 10^{5}$ | $1.79 \cdot 10^{6}$ |
| $k_{d5}$ | $[6.01 \cdot 10^{-4}, 6.01]$ | $7.85 \cdot 10^{-1}$ | 2.59 |
| $k_{d6}$ | $[6.01 \cdot 10^{-3}, 6.01 \cdot 10^{1}]$ | $2.37 \cdot 10^{1}$ | $1.43 \cdot 10^{1}$ |
| *Training hyper-parameters* | | | |
| learning rate | $[10^{-5}, 10^{-1}]$ | $2.57 \cdot 10^{-2}$ | $1.04 \cdot 10^{-2}$ |
| $\lambda$ | $\{10^{-3}, 10^{-2}, 10^{-1}, 1\}$ | ND | 0.01 |
| $k$ (segmentation) | $\{2, 3, 4, 5, 6, 7, 8, 9, 10\}$ | 2 | 8 |
| $\rho$ | $[10^{-3}, 10^{3}]$ | $5.24 \cdot 10^{-3}$ | $9.88 \cdot 10^{2}$ |

### S10 Cell apoptosis test case (first scenario, with $k_{d1}$ and $k_3$ fixed): identifiability analysis

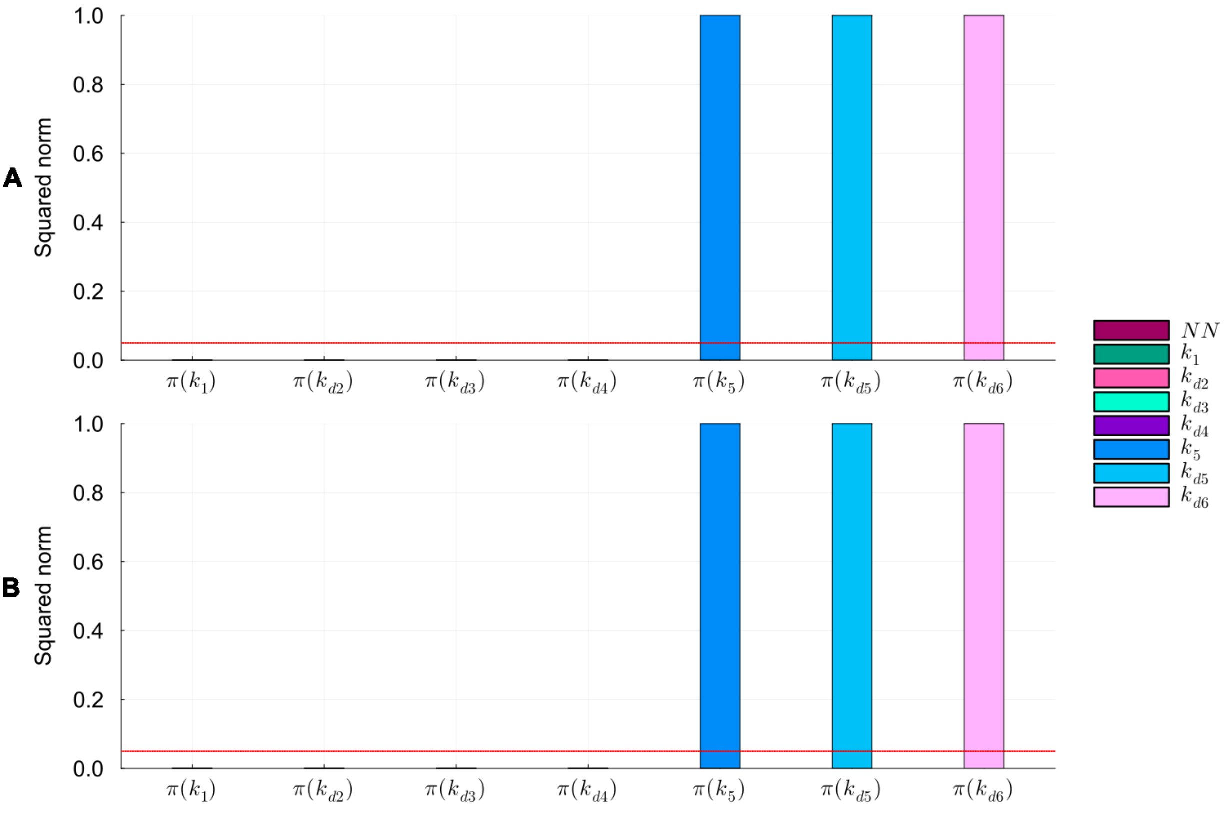


Figure S5: **Identifiability analysis of mechanistic parameters in the cell apoptosis test case (first scenario, with $k_{d1}$ and $k_3$ fixed).** Squared norm of the projections of the mechanistic parameters onto the null subspace of $H_\chi$ for the models trained on $DS_{0.00}$ and $DS_{0.05}$ (panel A and B respectively). The total height of the bar corresponds to the squared norm of the projection, while the different components of the projection are depicted in different colors. The red line indicates the threshold to determine the identifiability of the parameter (0.05).

#### S11 Cell apoptosis test case (first scenario, with $k_{d1}$ and $k_3$ fixed): predicted dynamics

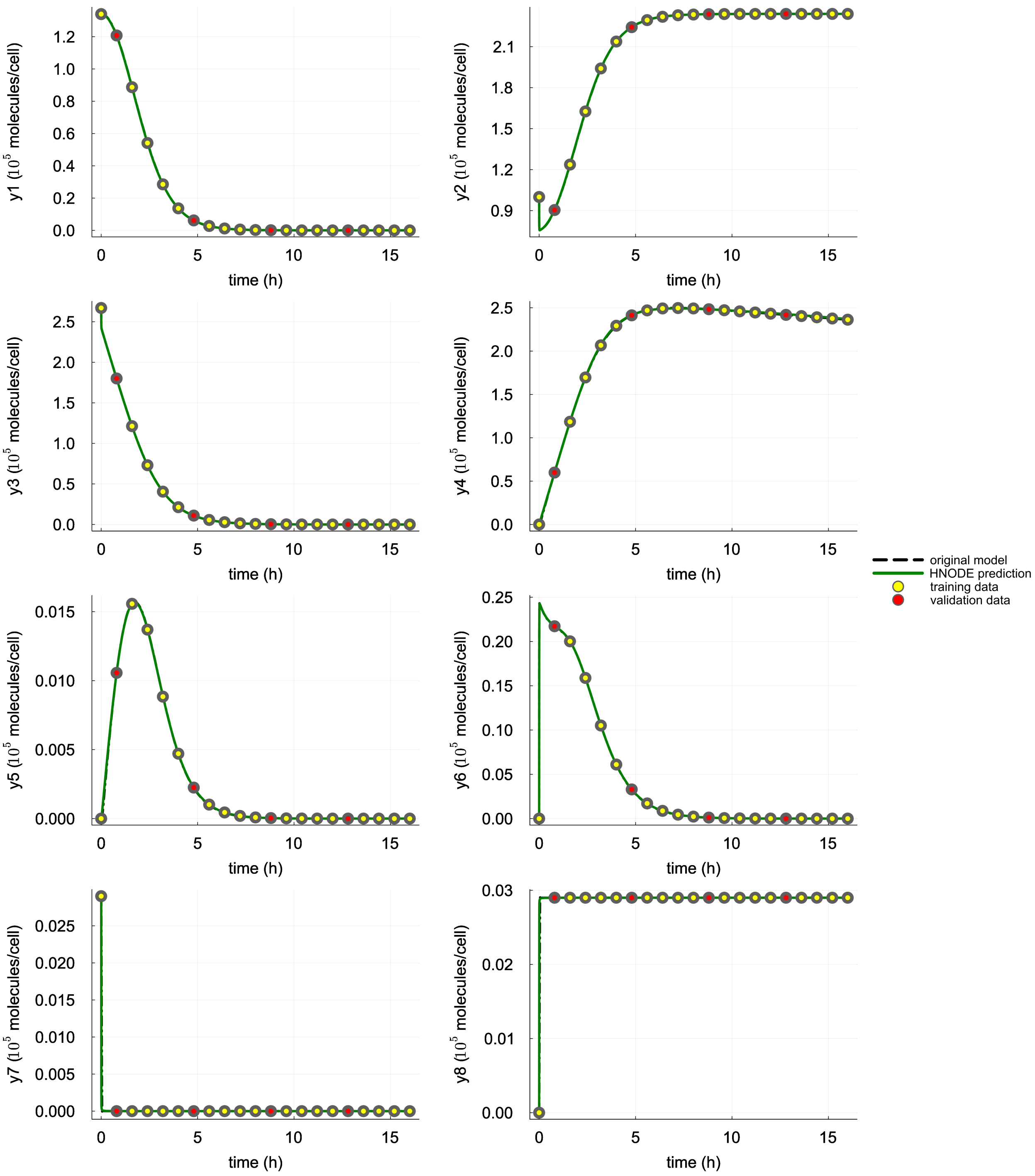


Figure S6: **Dynamics predicted by the HNODE model in the cell apoptosis test case (first scenario, with $k_{d1}$ and $k_3$ fixed).** The dynamics predicted by the HNODE model trained on $DS_{0.00}$ are compared with the original model. The points represent the observations of the system, divided into training and validation sets.

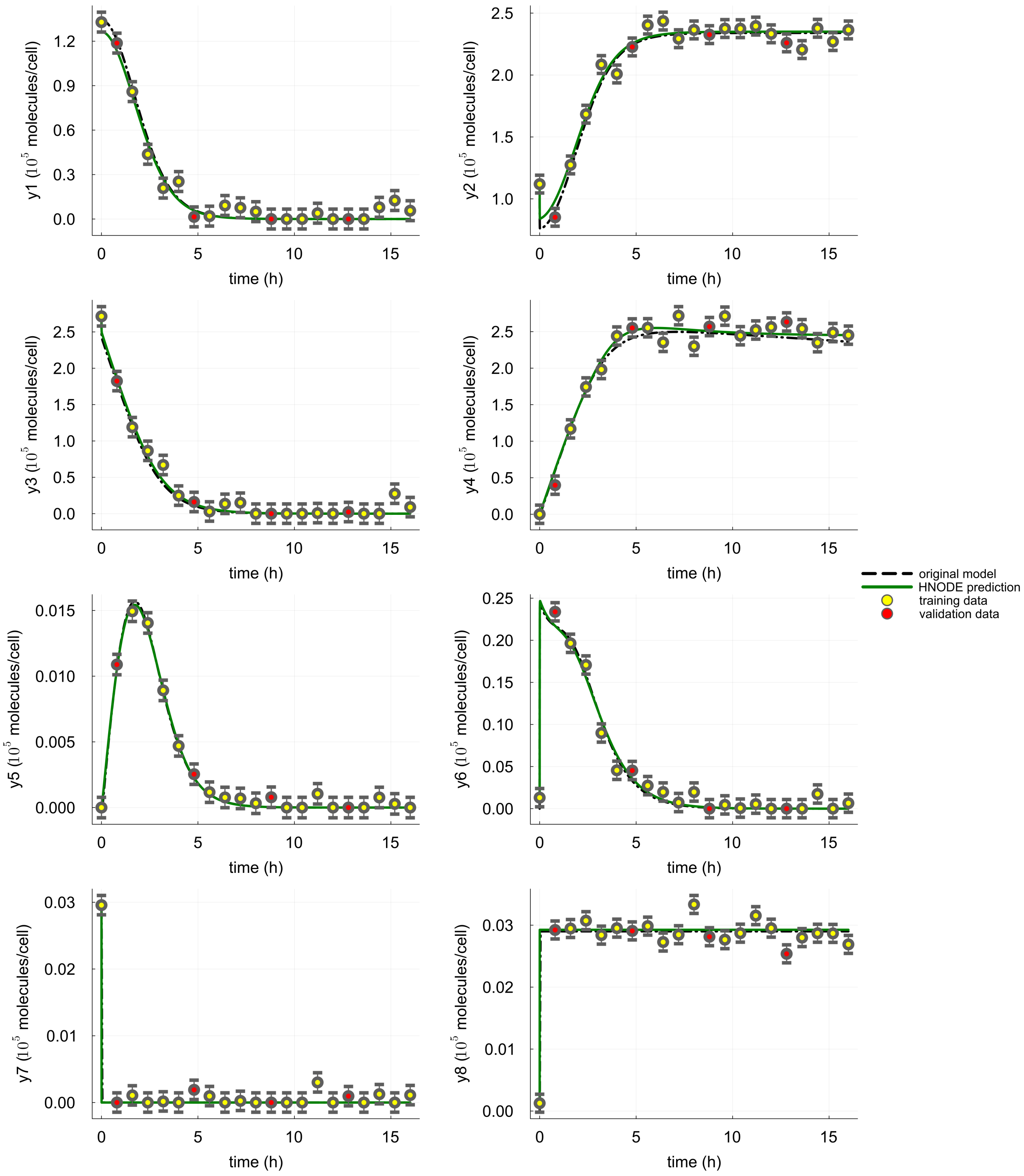


Figure S7: **Dynamics predicted by the HNODE model in the cell apoptosis test case (first scenario, with $k_{d1}$ and $k_3$ fixed).** The dynamics predicted by the HNODE model trained on $DS_{0.05}$ are compared with the original model. The points represent the observations of the system, divided into training and validation sets.

### S12 Cell apoptosis test case (second scenario): tuned hyper-parameters

Table S9: **Cell apoptosis test case (second scenario): tuned hyperparameters of the HNODE model.**

| Name | Search Space | Tuned value $DS_{0.00}$ | Tuned value $DS_{0.05}$ |
|---|---|---|---|
| *Neural network architecture* | | | |
| hidden layer number | $\{2, 3, 4, 5, 6\}$ | 2 | 5 |
| hidden layer width | $\{4, 8, 16, 32\}$ | 8 | 4 |
| *Starting values for mechanistic parameters* | | | |
| $k_1$ | $[9.61 \cdot 10^{-3}, 9.61 \cdot 10^{1}]$ | 6.73 | 2.08 |
| $k_{d2}$ | $[2.88 \cdot 10^{-1}, 2.88 \cdot 10^{3}]$ | $6.93 \cdot 10^{2}$ | $1.24 \cdot 10^{3}$ |
| $k_{d3}$ | $[1.80, 1.80 \cdot 10^{4}]$ | $8.82 \cdot 10^{2}$ | $1.59 \cdot 10^{4}$ |
| $k_{d4}$ | $[3.60 \cdot 10^{-2}, 3.60 \cdot 10^{2}]$ | $4.98 \cdot 10^{1}$ | $5.01 \cdot 10^{1}$ |
| $k_5$ | $[2.52 \cdot 10^{2}, 2.52 \cdot 10^{6}]$ | $2.39 \cdot 10^{6}$ | $2.28 \cdot 10^{6}$ |
| $k_{d5}$ | $[6.01 \cdot 10^{-4}, 6.01]$ | $8.98 \cdot 10^{-1}$ | 2.43 |
| $k_{d6}$ | $[6.01 \cdot 10^{-3}, 6.01 \cdot 10^{1}]$ | $2.70 \cdot 10^{1}$ | $3.14 \cdot 10^{1}$ |
| *Training hyper-parameters* | | | |
| learning rate | $[10^{-5}, 10^{-1}]$ | $4.14 \cdot 10^{-2}$ | $4.12 \cdot 10^{-3}$ |
| $\lambda$ | $\{10^{-3}, 10^{-2}, 10^{-1}, 1\}$ | ND | 0.001 |
| $k$ (segmentation) | $[2, 20]$ | 19 | 10 |
| $\rho$ | $[10^{-3}, 10^{3}]$ | $4.04 \cdot 10^{-3}$ | $4.97 \cdot 10^{-3}$ |

### S13 Cell apoptosis test case (second scenario): identifiability analysis

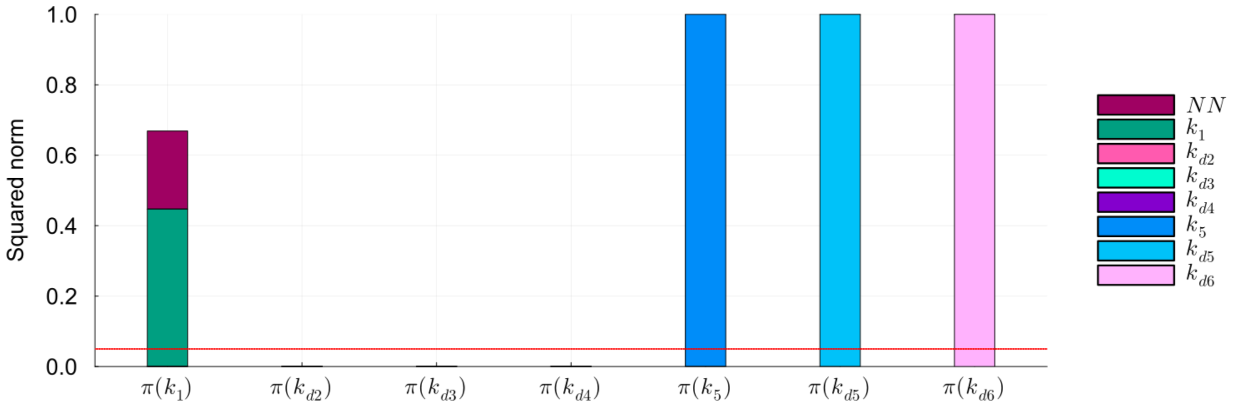


Figure S8: **Identifiability analysis of mechanistic parameters in the cell apoptosis test case, second scenario ($DS_{0.05}$).** Squared norm of the projections of the mechanistic parameters onto the null subspace of $H_\chi$ for the model trained on $DS_{0.05}$. The total height of the bar corresponds to the squared norm of the projection, while the different components of the projection are depicted in different colors. The red line indicates the threshold to determine the identifiability of the parameter (0.05).

### S14 Cell apoptosis test case (second scenario): predicted dynamics

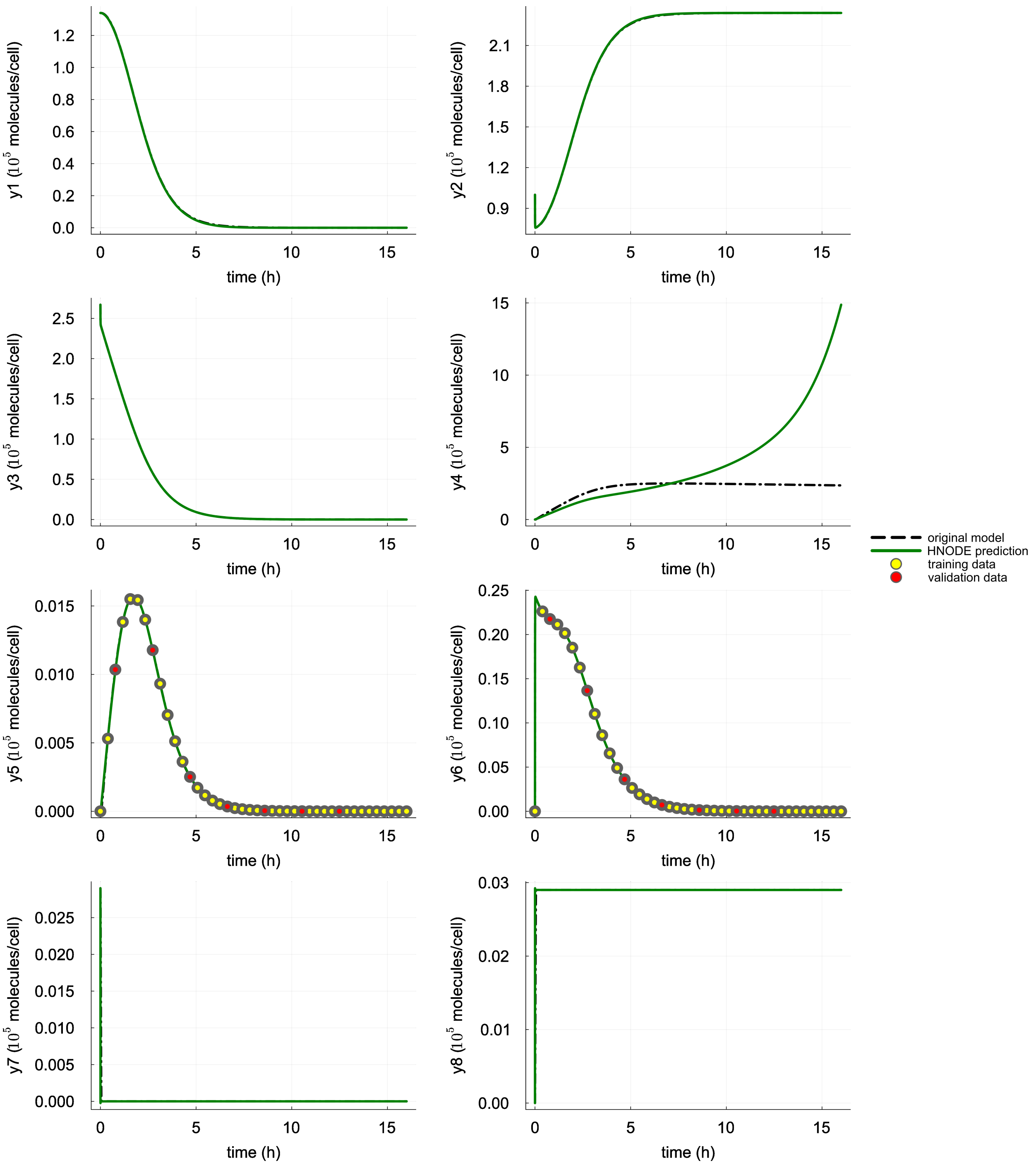


Figure S9: **Dynamics predicted by the HNODE model in the cell apoptosis test case, second scenario ($DS_{0.00}$).** The dynamics predicted by the HNODE model trained on $DS_{0.00}$ are compared with the original model. The points represent the observations of the system, divided into training and validation sets.

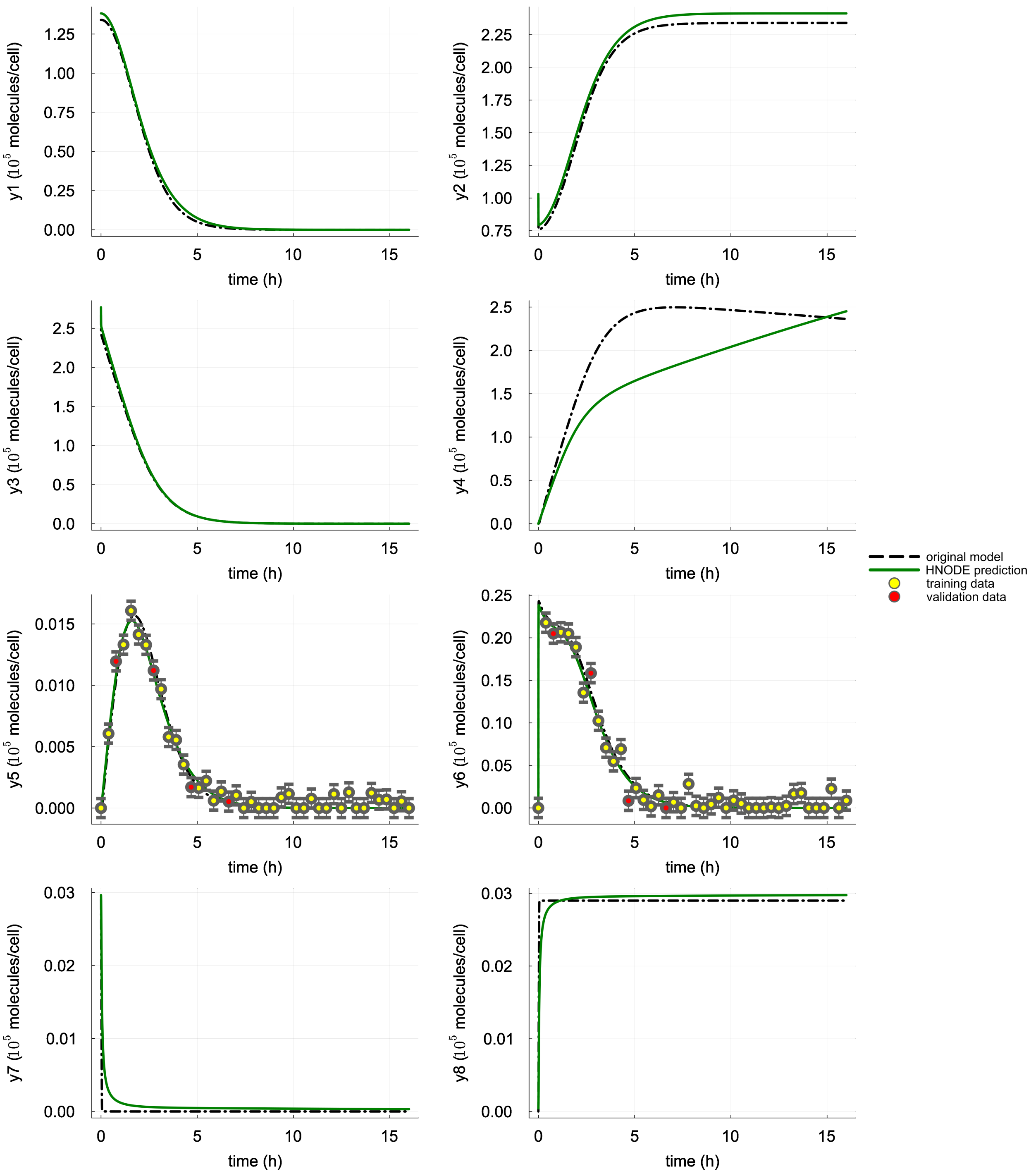


Figure S10: **Dynamics predicted by the HNODE model in the cell apoptosis test case, second scenario ($DS_{0.05}$).** The dynamics predicted by the HNODE model trained on $DS_{0.05}$ are compared with the original model. The points represent the observations of the system, divided into training and validation sets.

### S15 Cell apoptosis test case: identifiability analysis of the original mechanistic model with different combinations of observable variables

The local identifiability of parameters within the original mechanistic model for cell apoptosis was evaluated using the methodology proposed by the authors in [QM09]. This evaluation assumed the same observation time-points as the first scenario and explored various combinations of observable variables. Table S10 presents the identifiable parameters under different combinations of observable variables. Within the original model, $y_5$ and $y_6$ form a minimal subset of observable variables that maximizes the number of identifiable parameters.

Table S10: **Cell apoptosis: identifiability analysis of the original mechanistic model with various combinations of observable variables.**

| **Observable variables** | $k_1$ | $k_{d1}$ | $k_{d2}$ | $k_3$ | $k_{d3}$ | $k_{d4}$ | $k_5$ | $k_{d5}$ | $k_{d6}$ |
|---|---|---|---|---|---|---|---|---|---|
| y1, y2, y3, y4, y5, y6, y7, y8 | ✓ | | ✓ | | ✓ | ✓ | | | ✓ |
| y5, y6 | ✓ | | ✓ | | ✓ | ✓ | | | ✓ |
| y5 | ✓ | | ✓ | | | ✓ | | | |
| y6 | ✓ | | | | ✓ | | | | |

The symbol ✓ denotes that the corresponding parameter is classified as identifiable.

### S16 Cell apoptosis test case: impact of $\epsilon$ and $\delta$ on the identifiability analysis results

To assess the impact of varying hyperparameters $\epsilon$ and $\delta$ on the identifiability analysis, we conducted the analysis using different values for these parameters (Tables S11, S12, and S13). Our findings indicate that the results exhibit robustness against changes in $\epsilon$ and $\delta$.

Table S11: **Cell apoptosis test case (first scenario): parameter identifiability with different choices of $\epsilon$ and $\delta$.**

| | | | $\epsilon = 10^{-6}$ | | | $\epsilon = 10^{-5}$ | | | $\epsilon = 10^{-4}$ | |
|---|---|---|---|---|---|---|---|---|---|---|
| **Par.** | **Dataset** | $\delta = 0.03$ | $\delta = 0.05$ | $\delta = 0.07$ | $\delta = 0.03$ | $\delta = 0.05$ | $\delta = 0.07$ | $\delta = 0.05$ | $\delta = 0.07$ | |
| $k_1$ | $DS_{0.00}$ | | | | | | | | | |
| | $DS_{0.05}$ | | | | | | | | | |
| $k_{d1}$ | $DS_{0.00}$ | | | | | | | | | |
| | $DS_{0.05}$ | | | | | | | | | |
| $k_{d2}$ | $DS_{0.00}$ | ✓ | ✓ | ✓ | ✓ | ✓ | ✓ | ✓ | ✓ | ✓ |
| | $DS_{0.05}$ | ✓ | ✓ | ✓ | ✓ | ✓ | ✓ | ✓ | ✓ | ✓ |
| $k_3$ | $DS_{0.00}$ | | | | | | | | | |
| | $DS_{0.05}$ | | | | | | | | | |
| $k_{d3}$ | $DS_{0.00}$ | | | | | | | | | |
| | $DS_{0.05}$ | | | | | | | | | |
| $k_{d4}$ | $DS_{0.00}$ | ✓ | ✓ | ✓ | ✓ | ✓ | ✓ | ✓ | ✓ | ✓ |
| | $DS_{0.05}$ | ✓ | ✓ | ✓ | ✓ | ✓ | ✓ | ✓ | ✓ | ✓ |
| $k_5$ | $DS_{0.00}$ | | | | | | | | | |
| | $DS_{0.05}$ | | | | | | | | | |
| $k_{d5}$ | $DS_{0.00}$ | | | | | | | | | |
| | $DS_{0.05}$ | | | | | | | | | |
| $k_{d6}$ | $DS_{0.00}$ | | | | | | | | | |
| | $DS_{0.05}$ | | | | | | | | | |

The symbol ✓ denotes the parameters classified as identifiable with the $\epsilon$ and $\delta$ threshold specified in the header.

Table S12: **Cell apoptosis (first scenario, with $k_{d1}$ and $k_3$ fixed): parameter identifiability with different choices of $\epsilon$ and $\delta$.**

| | | | $\epsilon = 10^{-6}$ | | | $\epsilon = 10^{-5}$ | | $\epsilon = 10^{-4}$ | | |
|---|---|---|---|---|---|---|---|---|---|---|
| **Par.** | **Dataset** | $\delta = 0.03$ | $\delta = 0.05$ | $\delta = 0.07$ | $\delta = 0.03$ | $\delta = 0.05$ | $\delta = 0.07$ | $\delta = 0.05$ | $\delta = 0.07$ | |
| $k_1$ | $DS_{0.00}$ | ✓ | ✓ | ✓ | ✓ | ✓ | ✓ | ✓ | ✓ | ✓ |
| | $DS_{0.05}$ | ✓ | ✓ | ✓ | ✓ | ✓ | ✓ | ✓ | ✓ | ✓ |
| $k_{d2}$ | $DS_{0.00}$ | ✓ | ✓ | ✓ | ✓ | ✓ | ✓ | ✓ | ✓ | ✓ |
| | $DS_{0.05}$ | ✓ | ✓ | ✓ | ✓ | ✓ | ✓ | ✓ | ✓ | ✓ |
| $k_{d3}$ | $DS_{0.00}$ | ✓ | ✓ | ✓ | ✓ | ✓ | ✓ | ✓ | ✓ | ✓ |
| | $DS_{0.05}$ | ✓ | ✓ | ✓ | ✓ | ✓ | ✓ | ✓ | ✓ | ✓ |
| $k_{d4}$ | $DS_{0.00}$ | ✓ | ✓ | ✓ | ✓ | ✓ | ✓ | ✓ | ✓ | ✓ |
| | $DS_{0.05}$ | ✓ | ✓ | ✓ | ✓ | ✓ | ✓ | ✓ | ✓ | ✓ |
| $k_5$ | $DS_{0.00}$ | | | | | | | | | |
| | $DS_{0.05}$ | | | | | | | | | |
| $k_{d5}$ | $DS_{0.00}$ | | | | | | | | | |
| | $DS_{0.05}$ | | | | | | | | | |
| $k_{d6}$ | $DS_{0.00}$ | | | | | | | | | |
| | $DS_{0.05}$ | | | | | | | | | |

The symbol ✓ denotes the parameters classified as identifiable with the $\epsilon$ and $\delta$ threshold specified in the header.

Table S13: **Cell apoptosis (second scenario): parameter identifiability with different choices of $\epsilon$ and $\delta$.**

| | | | $\epsilon = 10^{-6}$ | | | $\epsilon = 10^{-5}$ | | $\epsilon = 10^{-4}$ | | |
|---|---|---|---|---|---|---|---|---|---|---|
| **Par.** | **Dataset** | $\delta = 0.03$ | $\delta = 0.05$ | $\delta = 0.07$ | $\delta = 0.03$ | $\delta = 0.05$ | $\delta = 0.07$ | $\delta = 0.05$ | $\delta = 0.07$ | |
| $k_1$ | $DS_{0.00}$ | | | | | | | | | |
| | $DS_{0.05}$ | | | | | | | | | |
| $k_{d2}$ | $DS_{0.00}$ | ✓ | ✓ | ✓ | ✓ | ✓ | ✓ | ✓ | ✓ | ✓ |
| | $DS_{0.05}$ | ✓ | ✓ | ✓ | ✓ | ✓ | ✓ | ✓ | ✓ | ✓ |
| $k_{d3}$ | $DS_{0.00}$ | ✓ | ✓ | ✓ | ✓ | ✓ | ✓ | ✓ | ✓ | ✓ |
| | $DS_{0.05}$ | ✓ | ✓ | ✓ | ✓ | ✓ | ✓ | ✓ | ✓ | ✓ |
| $k_{d4}$ | $DS_{0.00}$ | ✓ | ✓ | ✓ | ✓ | ✓ | ✓ | ✓ | ✓ | ✓ |
| | $DS_{0.05}$ | ✓ | ✓ | ✓ | ✓ | ✓ | ✓ | ✓ | ✓ | ✓ |
| $k_5$ | $DS_{0.00}$ | | | | | | | | | |
| | $DS_{0.05}$ | | | | | | | | | |
| $k_{d5}$ | $DS_{0.00}$ | | | | | | | | | |
| | $DS_{0.05}$ | | | | | | | | | |
| $k_{d6}$ | $DS_{0.00}$ | | | | | | | | | |
| | $DS_{0.05}$ | | | | | | | | | |

The symbol ✓ denotes the parameters classified as identifiable with the $\epsilon$ and $\delta$ threshold specified in the header.

#### S17 Yeast glycolysis test case: experimental setting

To simulate in silico observations, the Yeast glycolysis ODE system is solved numerically employing the *TRBDF2* solver from the *DifferentialEquations.jl* library. The solver settings are configured with an absolute tolerance of $10^{-7}$ and a relative tolerance of $10^{-6}$. The parameters and initial states, derived from [RCWH03], are detailed in Table S14 and S15.

Table S14: **Yeast glycolysis test case: parameters employed for generating the *in silico* dataset**

| **Parameter** | **Value** | **Unit** |
|---|---|---|
| $J_0$ | 2.5 | $\mathrm{mM} \cdot \mathrm{min}^{-1}$ |
| $k_1$ | 100.0 | $\mathrm{mM}^{-1} \cdot \mathrm{min}^{-1}$ |
| $k_2$ | 6.0 | $\mathrm{mM}^{-1} \cdot \mathrm{min}^{-1}$ |
| $k_3$ | 16.0 | $\mathrm{mM}^{-1} \cdot \mathrm{min}^{-1}$ |
| $k_4$ | 100.0 | $\mathrm{mM}^{-1} \cdot \mathrm{min}^{-1}$ |
| $k_5$ | 1.28 | $\mathrm{min}^{-1}$ |
| $k_6$ | 12.0 | $\mathrm{mM}^{-1} \cdot \mathrm{min}^{-1}$ |
| $k$ | 1.8 | $\mathrm{min}^{-1}$ |
| $\kappa$ | 13.0 | $\mathrm{min}^{-1}$ |
| $q$ | 4.0 | |
| $K_1$ | 0.52 | mM |
| $\psi$ | 0.1 | |
| $N$ | 1.0 | mM |
| $A$ | 4.0 | mM |

Table S15: **Yeast glycolysis test case: initial states employed for generating the *in silico* dataset**

| **Variable** | **Value** | **Unit** |
|---|---|---|
| $y_1$ | 1.6 | mM |
| $y_2$ | 2.16 | mM |
| $y_3$ | 0.2 | mM |
| $y_4$ | 0.35 | mM |
| $y_5$ | 0.3 | mM |
| $y_6$ | 2.67 | mM |
| $y_7$ | 0.1 | mM |

#### S18 Yeast glycolysis test case (first scenario): tuned hyper-parameters

Table S16: **Yeast glycolysis test case (first scenario): tuned hyperparameters of the HNODE model.**

| Name | Search Space | Tuned value $DS_{0.00}$ | Tuned value $DS_{0.05}$ |
|---|---|---|---|
| *Neural network architecture* | | | |
| hidden layer number | $\{2, 3, 4, 5, 6\}$ | 2 | 5 |
| hidden layer width | $\{4, 8, 16, 32\}$ | 16 | 16 |
| *Starting values for mechanistic parameters* | | | |
| $k_1$ | $[1.00 \cdot 10^1, 1.00 \cdot 10^3]$ | $5.70 \cdot 10^1$ | $1.15 \cdot 10^2$ |
| $K_1$ | $[5.20 \cdot 10^{-2}, 5.20]$ | $1.10 \cdot 10^{-1}$ | $6.39 \cdot 10^{-2}$ |
| $q$ | $[4.00 \cdot 10^{-1}, 4.00 \cdot 10^1]$ | 6.52 | 5.18 |
| $k_2$ | $[6.00 \cdot 10^{-1}, 6.00 \cdot 10^1]$ | $3.24 \cdot 10^1$ | 4.98 |
| $N$ | $[1.00 \cdot 10^{-1}, 1.00 \cdot 10^1]$ | 2.71 | $4.53 \cdot 10^{-1}$ |
| $k_6$ | $[1.20, 1.20 \cdot 10^2]$ | $2.68 \cdot 10^1$ | $1.71 \cdot 10^1$ |
| $k_3$ | $[1.60, 1.60 \cdot 10^2]$ | $1.21 \cdot 10^2$ | $4.92 \cdot 10^1$ |
| $A$ | $[4.00 \cdot 10^{-1}, 4.00 \cdot 10^1]$ | $2.13 \cdot 10^1$ | $1.82 \cdot 10^1$ |
| $k_4$ | $[1.00 \cdot 10^1, 1.00 \cdot 10^3]$ | $1.35 \cdot 10^2$ | $5.30 \cdot 10^2$ |
| $\kappa$ | $[1.30, 1.30 \cdot 10^2]$ | $3.76 \cdot 10^1$ | $2.02 \cdot 10^1$ |
| $k_5$ | $[1.28 \cdot 10^{-1}, 1.28 \cdot 10^1]$ | 3.81 | 7.05 |
| $\psi$ | $[1.00 \cdot 10^{-2}, 1.00]$ | $6.24 \cdot 10^{-1}$ | $1.76 \cdot 10^{-1}$ |
| $k$ | $[1.80 \cdot 10^{-1}, 1.80 \cdot 10^1]$ | 2.45 | 5.93 |
| *Training hyper-parameters* | | | |
| learning rate | $[10^{-5}, 10^{-1}]$ | $4.74 \cdot 10^{-2}$ | $1.07 \cdot 10^{-2}$ |
| $\lambda$ | $\{10^{-3}, 10^{-2}, 10^{-1}, 1\}$ | ND | 0.01 |
| $k$ (segmentation) | $\{2, 3, 4, 5, 6, 7, 8, 9, 10\}$ | 10 | 5 |
| $\rho$ | $[10^{-3}, 10^3]$ | $2.03 \cdot 10^{-2}$ | $5.43 \cdot 10^{-1}$ |

#### S19 Yeast glycolysis test case (first scenario): predicted dynamics

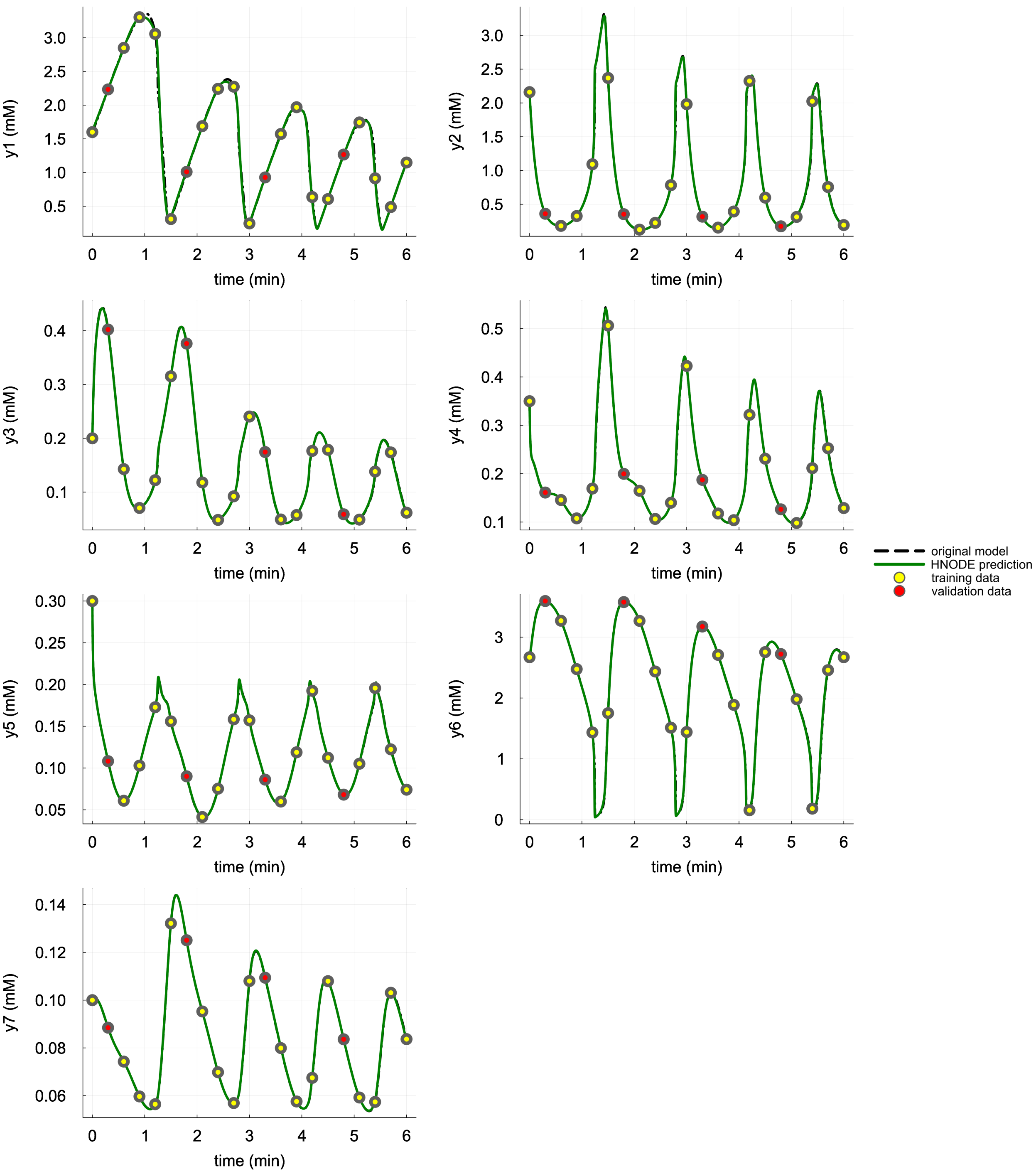


Figure S11: **Dynamics predicted by the HNODE model in the yeast glycolysis test case, first scenario ($DS_{0.00}$).** The dynamics predicted by the HNODE model trained on $DS_{0.00}$ are compared with the original model. The points represent the observations of the system, divided into training and validation sets.

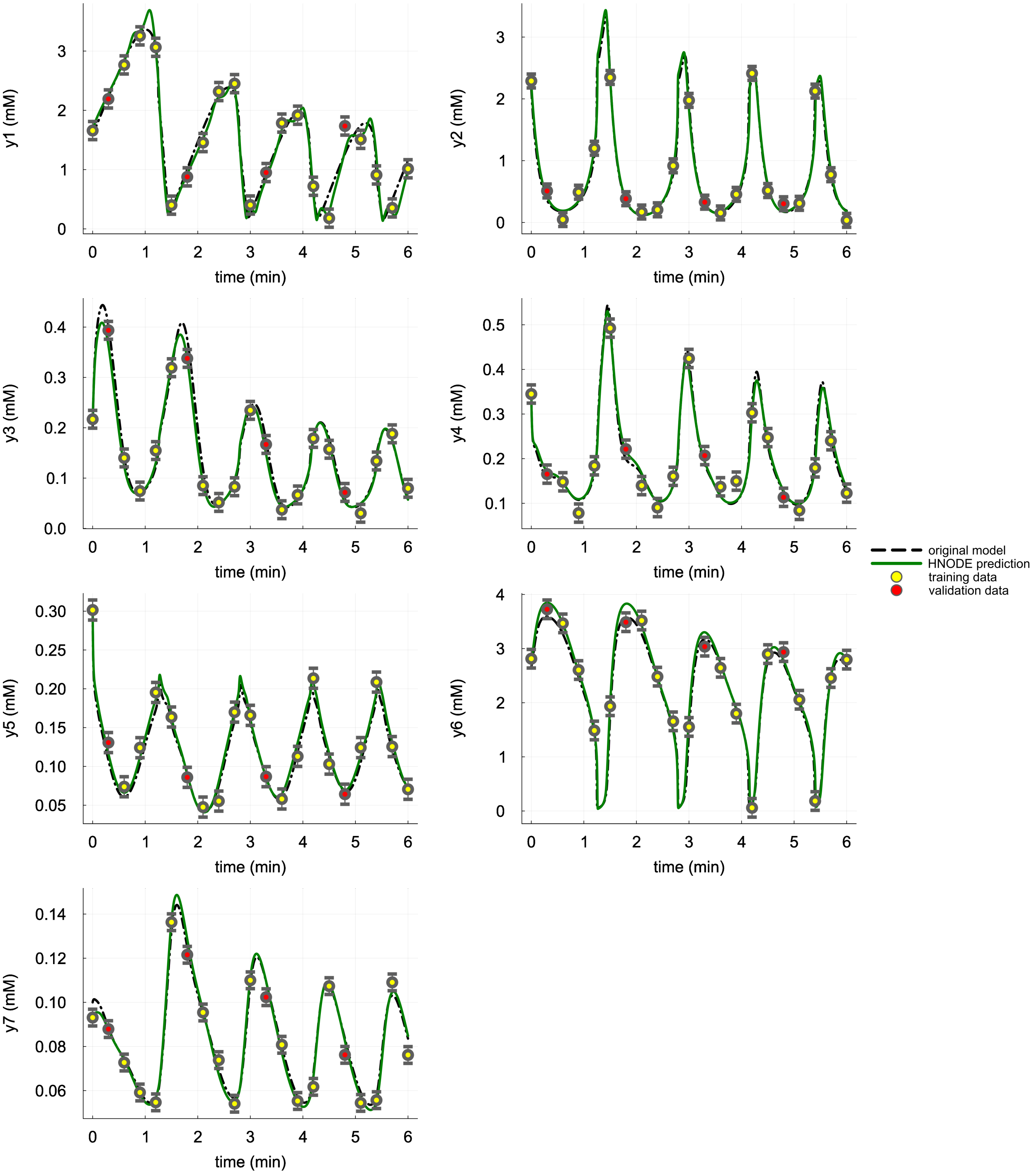


Figure S12: **Dynamics predicted by the HNODE model in the yeast glycolysis test case, first scenario ($DS_{0.05}$).** The dynamics predicted by the HNODE model trained on $DS_{0.05}$ are compared with the original model. The points represent the observations of the system, divided into training and validation sets.

#### S20 Yeast glycolysis test case (first scenario): identifiability analysis

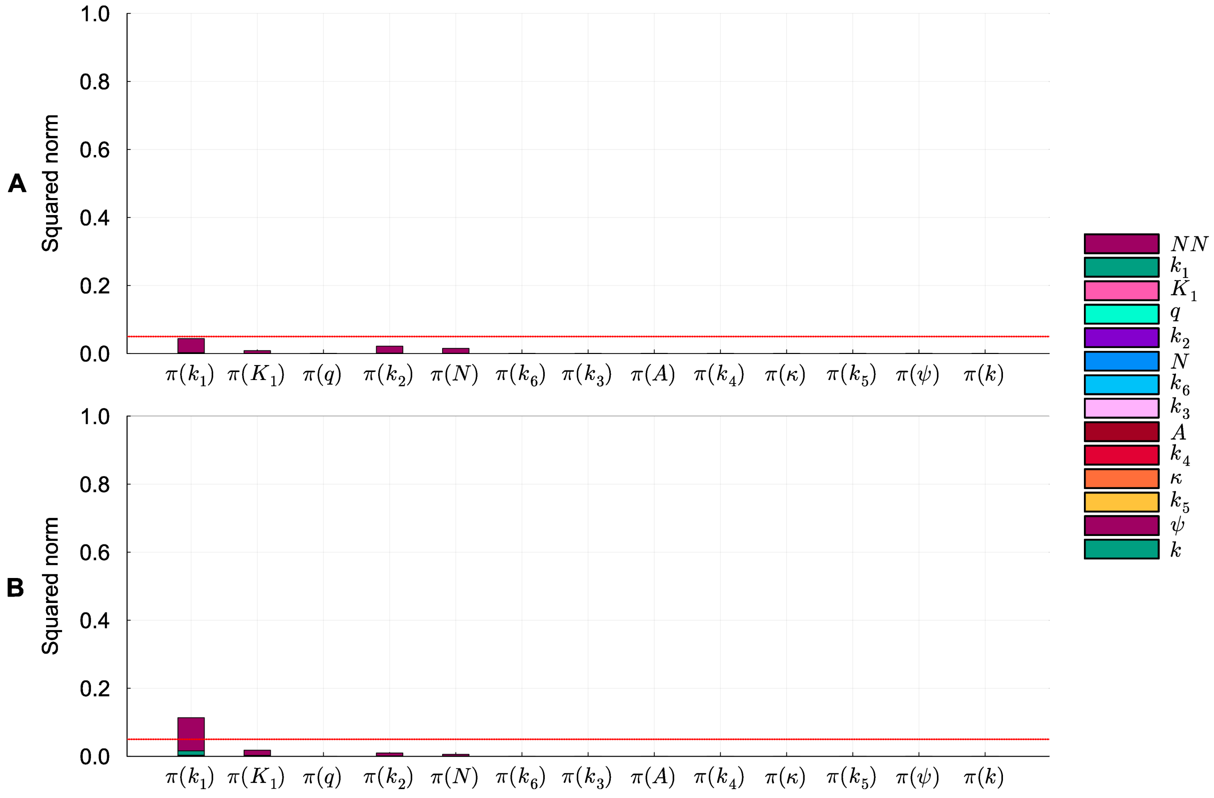


Figure S13: **Identifiability analysis of mechanistic parameters in the yeast glycolysis test case, first scenario.** Squared norm of the projections of the mechanistic parameters onto the null subspace of $H_\chi$ for the models trained on $DS_{0.00}$ and $DS_{0.05}$ (panel A and B respectively). The total height of the bar corresponds to the squared norm of the projection, while the different components of the projection are depicted in different colors. The red line indicates the threshold to determine the identifiability of the parameter (0.05).

### S21 Yeast glycolysis test case (second scenario): tuned hyperparameters

Table S17: **Yeast glycolysis test case (second scenario): tuned hyperparameters of the HNODE model.**

| Name | Search Space | Tuned value $DS_{0.00}$ | Tuned value $DS_{0.05}$ |
|---|---|---|---|
| *Neural network architecture* | | | |
| hidden layer number | $\{2,3,4,5,6\}$ | 3 | 2 |
| hidden layer width | $\{4,8,16,32\}$ | 16 | 16 |
| *Starting values for mechanistic parameters* | | | |
| $k_1$ | $[1.00\cdot10^1, 1.00\cdot10^3]$ | $8.96\cdot10^1$ | $6.50\cdot10^1$ |
| $K_1$ | $[5.20\cdot10^{-2}, 5.20]$ | $9.09\cdot10^{-2}$ | $2.35\cdot10^{-1}$ |
| $q$ | $[4.00\cdot10^{-1}, 4.00\cdot10^1]$ | 5.74 | 3.34 |
| $k_2$ | $[6.00\cdot10^{-1}, 6.00\cdot10^1]$ | 3.10 | 1.83 |
| $N$ | $[1.00\cdot10^{-1}, 1.00\cdot10^1]$ | 3.31 | 1.87 |
| $k_6$ | $[1.20, 1.20\cdot10^2]$ | 1.39 | $5.53\cdot10^1$ |
| $k_3$ | $[1.60, 1.60\cdot10^2]$ | 4.54 | $2.82\cdot10^1$ |
| $A$ | $[4.00\cdot10^{-1}, 4.00\cdot10^1]$ | $1.94\cdot10^1$ | $1.64\cdot10^1$ |
| $k_4$ | $[1.00\cdot10^1, 1.00\cdot10^3]$ | $8.66\cdot10^2$ | $5.05\cdot10^2$ |
| $\kappa$ | $[1.30, 1.30\cdot10^2]$ | $7.02\cdot10^1$ | $1.10\cdot10^1$ |
| $k_5$ | $[1.28\cdot10^{-1}, 1.28\cdot10^1]$ | 2.09 | 1.40 |
| $\psi$ | $[1.00\cdot10^{-2}, 1.00]$ | $7.97\cdot10^{-1}$ | $2.09\cdot10^{-1}$ |
| $k$ | $[1.80\cdot10^{-1}, 1.80\cdot10^1]$ | $1.74\cdot10^1$ | 8.71 |
| *Training hyper-parameters* | | | |
| learning rate | $[10^{-5}, 10^{-1}]$ | $2.10\cdot10^{-2}$ | $7.90\cdot10^{-2}$ |
| $\lambda$ | $\{10^{-3}, 10^{-2}, 10^{-1}, 1\}$ | ND | 0.1 |
| $k$ (segmentation) | $[2, 20]$ | 19 | 9 |
| $\rho$ | $[10^{-3}, 10^3]$ | $1.94\cdot10^{-3}$ | $7.05\cdot10^{-1}$ |

### S22 Yeast glycolysis test case (second scenario): predicted dynamics

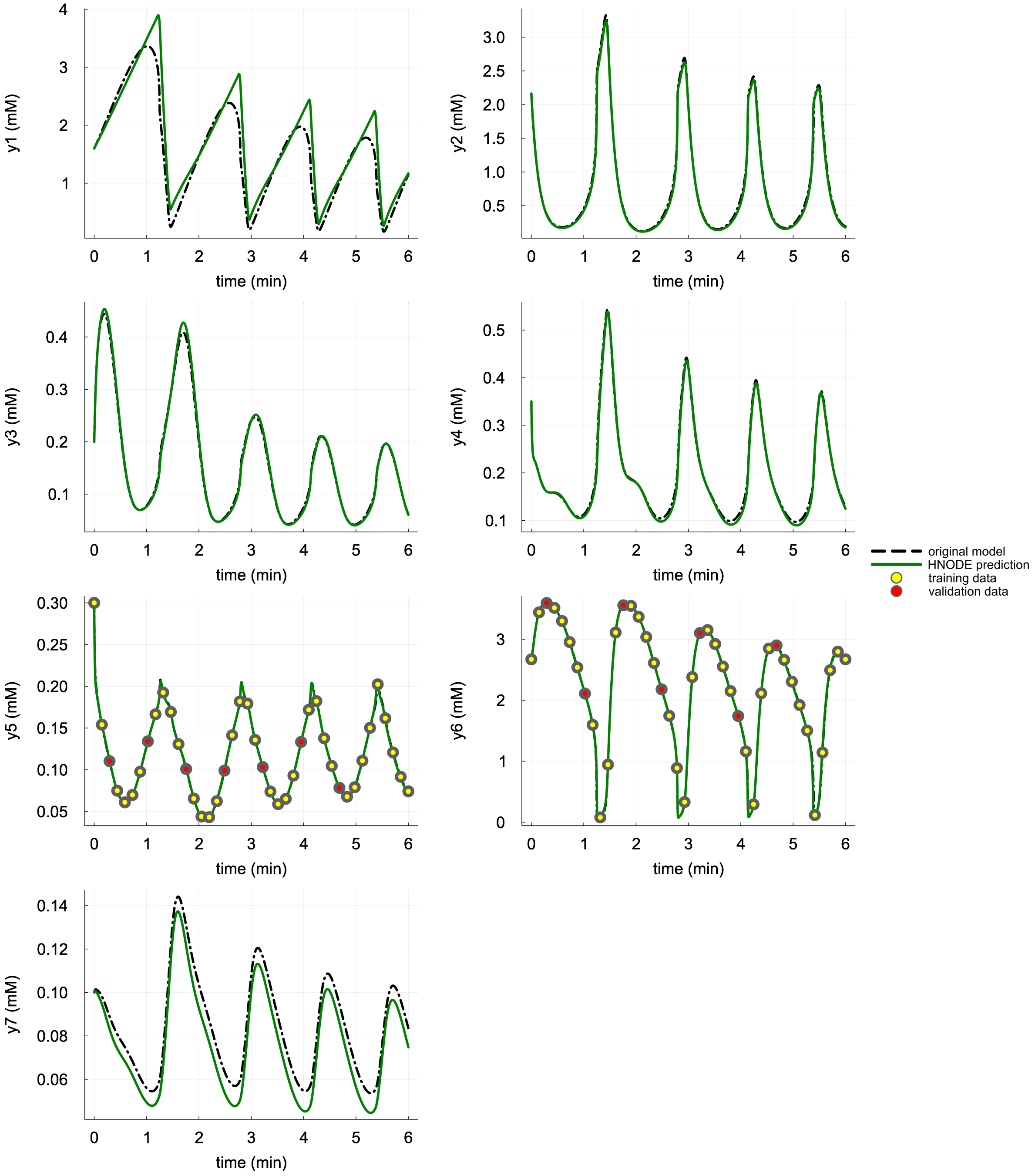


Figure S14: **Dynamics predicted by the HNODE model in the yeast glycolysis test case, second scenario ($DS_{0.00}$).** The dynamics predicted by the HNODE model trained on $DS_{0.00}$ are compared with the original model. The points represent the observations of the system, divided into training and validation sets.

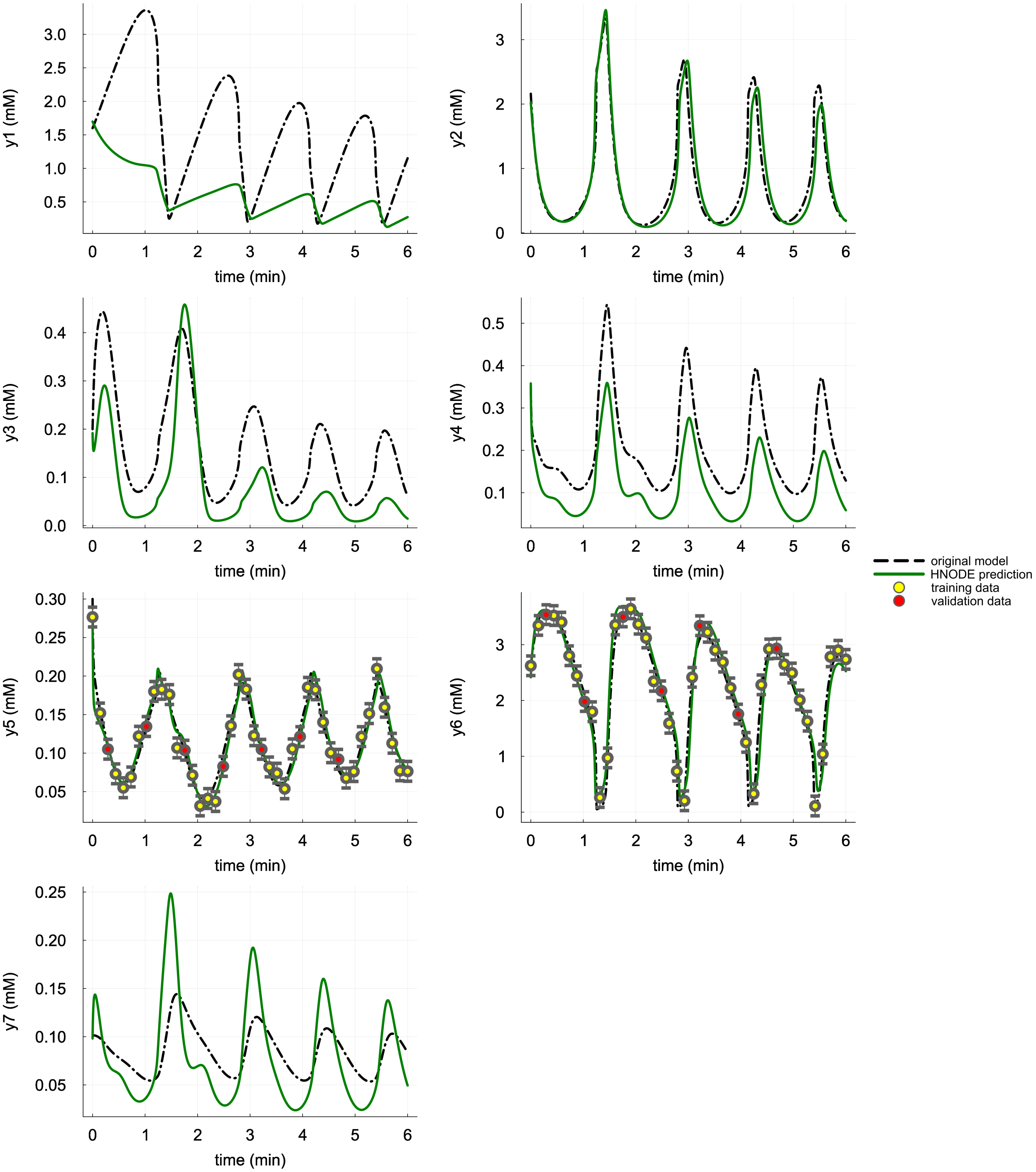


Figure S15: **Dynamics predicted by the HNODE model in the yeast glycolysis test case, second scenario ($DS_{0.05}$).** The dynamics predicted by the HNODE model trained on $DS_{0.05}$ are compared with the original model. The points represent the observations of the system, divided into training and validation sets.

### S23 Yeast glycolysis test case: parameter estimation with the original mechanistic model on the dataset $DS_{0.05}$

To distinguish between the effects of the lack of mechanistic knowledge and the noise in the datasets on parameter estimation, we conducted parameter estimation of the mechanistic model using the dataset $DS_{0.05}$. The estimation was carried out by minimizing the quadratic cost, weighted with uncertainty, assuming 21 observation time-points. Optimization was performed using the L-BFGS algorithm until convergence, with parameters initialized to their literature values. The results are presented in Table S18.

Table S18: **Yeast glycolysis test case: parameters estimated on the dataset $DS_{0.05}$ using the original model.**

| **Par.** | **Target** | **Estimate** | **Relative Error** |
|---|---|---|---|
| $J_0$ | 2.50 | 2.50 | 0.14% |
| $k_1$ | $1.00 \cdot 10^2$ | $1.29 \cdot 10^2$ | 29.23% |
| $K_1$ | $5.20 \cdot 10^{-1}$ | $5.01 \cdot 10^{-1}$ | 3.65% |
| $q$ | 4.00 | 4.03 | 0.82% |
| $k_2$ | 6.00 | 1.82 | 69.69% |
| $N$ | 1.00 | 2.88 | 188.19% |
| $k_6$ | $1.20 \cdot 10^1$ | $1.11 \cdot 10^1$ | 7.76% |
| $k_3$ | $1.60 \cdot 10^1$ | $1.53 \cdot 10^1$ | 4.36% |
| $A$ | 4.00 | 4.11 | 2.75% |
| $k_4$ | $1.00 \cdot 10^2$ | $9.34 \cdot 10^1$ | 6.59% |
| $\kappa$ | $1.30 \cdot 10^1$ | $1.31 \cdot 10^1$ | 0.88% |
| $k_5$ | 1.28 | 1.25 | 2.69% |
| $\psi$ | $1.00 \cdot 10^{-1}$ | $1.03 \cdot 10^{-1}$ | 2.93% |
| $k$ | 1.80 | 1.90 | 5.44% |
| | | **Mean** | 23.22% |
| | | **Max** | 188.19% |

### S24 Yeast glycolysis test case: impact of $\epsilon$ and $\delta$ on the identifiability analysis results

To assess the impact of varying hyperparameters $\epsilon$ and $\delta$ on the identifiability analysis, we conducted the analysis using different values for these parameters. In the first scenario (Table S19), the results demonstrate overall robustness, except for parameters $k_1$, $K_1$, $k_2$, and $N$, which appear to be influenced by compensatory mechanisms, particularly noticeable when considering $\epsilon = 10^{-4}$. In the second scenario (S20), there are differences between the HNODE model trained on $DS_{0.00}$ and $DS_{0.05}$. However, overall, only the results related to parameters $q$, $A$, $k_2$, and $k_5$ appear to maintain robustness against variations in $\epsilon$. It is important to note that even in the original model, there exist sloppy directions where the model behavior changes only slightly compared to other directions. Choosing a higher value for $\epsilon$ could potentially include these sloppy, albeit not null-change, directions into the null subspace of $H_\chi$, resulting in an overly conservative assessment of parameter identifiability.

Table S19: **Yeast glycolysis, first scenario - parameter identifiability with different choices of $\epsilon$ and $\delta$.**

| | | | $\epsilon = 10^{-6}$ | | | $\epsilon = 10^{-5}$ | | | $\epsilon = 10^{-4}$ | |
|---|---|---|---|---|---|---|---|---|---|---|
| **Par.** | **Dataset** | $\delta = 0.03$ | $\delta = 0.05$ | $\delta = 0.07$ | $\delta = 0.03$ | $\delta = 0.05$ | $\delta = 0.07$ | $\delta = 0.05$ | $\delta = 0.07$ | |
| $k_1$ | $DS_{0.00}$ | ✓ | ✓ | ✓ | | ✓ | ✓ | | | |
| | $DS_{0.05}$ | ✓ | ✓ | ✓ | | | | | | |
| $K_1$ | $DS_{0.00}$ | ✓ | ✓ | ✓ | ✓ | ✓ | ✓ | | | |
| | $DS_{0.05}$ | ✓ | ✓ | ✓ | ✓ | ✓ | ✓ | | | |
| $q$ | $DS_{0.00}$ | ✓ | ✓ | ✓ | ✓ | ✓ | ✓ | ✓ | ✓ | ✓ |
| | $DS_{0.05}$ | ✓ | ✓ | ✓ | ✓ | ✓ | ✓ | ✓ | ✓ | ✓ |
| $k_2$ | $DS_{0.00}$ | ✓ | ✓ | ✓ | ✓ | ✓ | ✓ | | | |
| | $DS_{0.05}$ | ✓ | ✓ | ✓ | ✓ | ✓ | ✓ | | | |
| $N$ | $DS_{0.00}$ | ✓ | ✓ | ✓ | ✓ | ✓ | ✓ | | | |
| | $DS_{0.05}$ | ✓ | ✓ | ✓ | ✓ | ✓ | ✓ | | | |
| $k_6$ | $DS_{0.00}$ | ✓ | ✓ | ✓ | ✓ | ✓ | ✓ | ✓ | ✓ | ✓ |
| | $DS_{0.05}$ | ✓ | ✓ | ✓ | ✓ | ✓ | ✓ | ✓ | ✓ | ✓ |
| $k_3$ | $DS_{0.00}$ | ✓ | ✓ | ✓ | ✓ | ✓ | ✓ | ✓ | ✓ | ✓ |
| | $DS_{0.05}$ | ✓ | ✓ | ✓ | ✓ | ✓ | ✓ | ✓ | ✓ | ✓ |
| $A$ | $DS_{0.00}$ | ✓ | ✓ | ✓ | ✓ | ✓ | ✓ | ✓ | ✓ | ✓ |
| | $DS_{0.05}$ | ✓ | ✓ | ✓ | ✓ | ✓ | ✓ | ✓ | ✓ | ✓ |
| $k_4$ | $DS_{0.00}$ | ✓ | ✓ | ✓ | ✓ | ✓ | ✓ | ✓ | ✓ | ✓ |
| | $DS_{0.05}$ | ✓ | ✓ | ✓ | ✓ | ✓ | ✓ | ✓ | ✓ | ✓ |
| $\kappa$ | $DS_{0.00}$ | ✓ | ✓ | ✓ | ✓ | ✓ | ✓ | ✓ | ✓ | ✓ |
| | $DS_{0.05}$ | ✓ | ✓ | ✓ | ✓ | ✓ | ✓ | ✓ | ✓ | ✓ |
| $k_5$ | $DS_{0.00}$ | ✓ | ✓ | ✓ | ✓ | ✓ | ✓ | ✓ | ✓ | ✓ |
| | $DS_{0.05}$ | ✓ | ✓ | ✓ | ✓ | ✓ | ✓ | ✓ | ✓ | ✓ |
| $\psi$ | $DS_{0.00}$ | ✓ | ✓ | ✓ | ✓ | ✓ | ✓ | ✓ | ✓ | ✓ |
| | $DS_{0.05}$ | ✓ | ✓ | ✓ | ✓ | ✓ | ✓ | | ✓ | ✓ |
| $k$ | $DS_{0.00}$ | ✓ | ✓ | ✓ | ✓ | ✓ | ✓ | ✓ | ✓ | ✓ |
| | $DS_{0.05}$ | ✓ | ✓ | ✓ | ✓ | ✓ | ✓ | ✓ | ✓ | ✓ |

The symbol ✓ denotes the parameters classified as identifiable with the $\epsilon$ and $\delta$ threshold specified in the header.

Table S20: **Yeast glycolysis, second scenario - parameter identifiability with different choices of $\epsilon$ and $\delta$.**

| | | | $\epsilon = 10^{-6}$ | | | $\epsilon = 10^{-5}$ | | $\epsilon = 10^{-4}$ | | |
|---|---|---|---|---|---|---|---|---|---|---|
| **Par.** | **Dataset** | $\delta = 0.03$ | $\delta = 0.05$ | $\delta = 0.07$ | $\delta = 0.03$ | $\delta = 0.05$ | $\delta = 0.07$ | $\delta = 0.05$ | $\delta = 0.07$ | |
| $k_1$ | $DS_{0.00}$ | ✓ | ✓ | ✓ | | | | | | |
| | $DS_{0.05}$ | ✓ | ✓ | ✓ | | | | | | |
| $K_1$ | $DS_{0.00}$ | ✓ | ✓ | ✓ | ✓ | ✓ | ✓ | | | |
| | $DS_{0.05}$ | ✓ | ✓ | ✓ | | | ✓ | | | |
| $q$ | $DS_{0.00}$ | ✓ | ✓ | ✓ | ✓ | ✓ | ✓ | ✓ | ✓ | ✓ |
| | $DS_{0.05}$ | ✓ | ✓ | ✓ | ✓ | ✓ | ✓ | | ✓ | ✓ |
| $k_2$ | $DS_{0.00}$ | | | | | | | | | |
| | $DS_{0.05}$ | | | | | | | | | |
| $N$ | $DS_{0.00}$ | | ✓ | ✓ | | | | | | |
| | $DS_{0.05}$ | | | | | | | | | |
| $k_6$ | $DS_{0.00}$ | ✓ | ✓ | ✓ | ✓ | ✓ | ✓ | | | |
| | $DS_{0.05}$ | ✓ | ✓ | ✓ | ✓ | ✓ | ✓ | | | |
| $k_3$ | $DS_{0.00}$ | ✓ | ✓ | ✓ | | ✓ | ✓ | | | |
| | $DS_{0.05}$ | ✓ | ✓ | ✓ | | | | | | |
| $A$ | $DS_{0.00}$ | ✓ | ✓ | ✓ | ✓ | ✓ | ✓ | ✓ | ✓ | ✓ |
| | $DS_{0.05}$ | ✓ | ✓ | ✓ | ✓ | ✓ | ✓ | ✓ | ✓ | ✓ |
| $k_4$ | $DS_{0.00}$ | | ✓ | ✓ | | | ✓ | | | |
| | $DS_{0.05}$ | ✓ | ✓ | ✓ | | | | | | |
| $\kappa$ | $DS_{0.00}$ | | | | | | | | | |
| | $DS_{0.05}$ | ✓ | ✓ | ✓ | | | | | | |
| $k_5$ | $DS_{0.00}$ | ✓ | ✓ | ✓ | ✓ | ✓ | ✓ | ✓ | ✓ | ✓ |
| | $DS_{0.05}$ | ✓ | ✓ | ✓ | ✓ | ✓ | ✓ | | ✓ | ✓ |
| $\psi$ | $DS_{0.00}$ | | | | | | | | | |
| | $DS_{0.05}$ | ✓ | ✓ | ✓ | | | | | | |
| $k$ | $DS_{0.00}$ | | | | | | | | | |
| | $DS_{0.05}$ | ✓ | ✓ | ✓ | | | | | | |

The symbol ✓ denotes the parameters classified as identifiable with the $\epsilon$ and $\delta$ threshold specified in the header.

### References


[AHSL06] Bree B Aldridge, George Haller, Peter K Sorger, and Douglas A Lauffenburger. Direct lyapunov exponent analysis enables parametric study of transient signalling governing cell behaviour. *IEE Proceedings-Systems Biology*, 153(6):425–432, 2006.

[QM09] Tom Quaiser and Martin Mönnigmann. Systematic identifiability testing for unambiguous mechanistic modeling–application to jak-stat, map kinase, and nf-$\kappa$ b signaling pathway models. *BMC systems biology*, 3(1):1–21, 2009.

[RCWH03] Peter Ruoff, Melinda K Christensen, Jana Wolf, and Reinhart Heinrich. Temperature dependency and temperature compensation in a model of yeast glycolytic oscillations. *Biophysical chemistry*, 106(2):179–192, 2003.

[RMM+20] Christopher Rackauckas, Yingbo Ma, Julius Martensen, Collin Warner, Kirill Zubov, Rohit Supekar, Dominic Skinner, Ali Ramadhan, and Alan Edelman. Universal differential equations for scientific machine learning. *arXiv preprint arXiv:2001.04385*, 2020.
